# Deep generative modeling of sample-level heterogeneity in single-cell genomics

**DOI:** 10.1101/2022.10.04.510898

**Authors:** Pierre Boyeau, Justin Hong, Adam Gayoso, Martin Kim, José L. McFaline-Figueroa, Michael I. Jordan, Elham Azizi, Can Ergen, Nir Yosef

**Author notes:** These authors contributed equally (Pierre Boyeau, Justin Hong). These authors contributed equally (Can Ergen, Nir Yosef). Correspondence, nir.

## Abstract

The field of single-cell genomics is now observing a marked increase in the prevalence of cohort-level studies that include hundreds of samples and feature complex designs. These data have tremendous potential for discovering how sample or tissue-level phenotypes relate to cellular and molecular composition. However, current analyses are based on simplified representations of these data by averaging information across cells. We present MrVI, a deep generative model designed to realize the potential of cohort studies at the single-cell level. MrVI tackles two fundamental and intertwined problems: stratifying samples into groups and evaluating the cellular and molecular differences between groups, both without requiring *a priori* grouping of cells into types or states. Due to its single-cell perspective, MrVI is able to detect clinically relevant stratifications of patients in COVID-19 and inflammatory bowel disease (IBD) cohorts that are only manifested in certain cellular subsets, thus enabling new discoveries that would otherwise be overlooked. Similarly, we demonstrate that MrVI can de-novo identify groups of small molecules with similar biochemical properties and evaluate their effects on cellular composition and gene expression in large-scale perturbation studies. MrVI is available as open source at `scvi-tools.org`.

## 1 Introduction

Over the past two decades, the use of functional genomics in large-scale, many-sample studies has been instrumental in advancing our understanding of how clinical, genetic, and environmental properties are manifested at the cellular and molecular levels [1, 2]. These studies now benefit from a potentially transformative increase in quality and resolution thanks to the maturation of large-scale single-cell genomics, which provides access to detailed information about the cellular and molecular composition of hundreds of samples [3, 4, 5, 6, 7, 8, 9]. Realizing the potential of large-scale single-cell genomics, however, requires rethinking the analysis strategy. While early on most studies relied on small numbers of samples and focused on variation between cells, the emergence of large-scale single-cell genomics now opens the way for a more in-depth understanding of the variation between samples.

There are at least two fundamental tasks in such a sample-level analysis. The first, which we refer to as *exploratory* analysis, is to divide the samples into groups based on their respective cellular and molecular properties. This idea of *de novo* grouping has seen powerful applications in clinical studies that use functional genomics to enable more precise prognoses and treatment planning [10, 11]. As a prominent example, pan-cancer analysis with functional genomics revealed that, in surprisingly many cases, cancer patients are more effectively classified using their molecular data rather than histopathology [12]. The second task is to conduct *comparative* analysis, i.e., identify cellular and molecular features that differ between pre-defined groups of samples (e.g., cases vs controls). In bulk-level studies, this was usually done in the form of differential expression (DE) for detecting gene expression programs that are associated with conditions of interest [13]. The advent of single-cell genomics also popularized differential abundance (DA) as another form of comparison, one which seeks to discover cell states that are disproportionately abundant in a certain group of samples [14].

Current approaches for these two closely related problems suffer from limitations that preclude them from taking full advantage of the resolution afforded by single-cell genomics. Starting with exploratory analysis, a common approach for quantifying the distances between samples is to first organize the cells into groups (representing types or states) and then evaluate the differences in the frequency of each group [5, 7, 15, 16, 17]. This approach, however, may oversimplify the task by reducing the rich information we have about each sample. Furthermore, it hinges on the effective clustering of the cells (so as to represent distinct cell states), which is often complicated by the need to set an ample level of resolution, distinguish between closely related states, and harmonize different samples or datasets. Finally, this approach can miss critical effects that may only manifest in particular subsets of cells (As we later demonstrate, using cohorts of IBD and COVID patients). Similar issues also emerge in comparative analyses. Most of the current applications of DE and DA rely on *a priori* clustering of cells. It is possible, however, that DE programs span few or parts of the *a priori* defined cell subsets, thus less likely to be detected. Similarly, differentially abundant sub-populations may not clearly correspond to any annotated subset, which again limits the ability to detect them. Finally, even with access to high-quality annotation of the cells, it may still be that comparative analyses of different partitions (e.g., comparing between sexes or between age groups) are best reflected by different cell clustering schemes [18].

To mitigate these issues, a recent line of work focused on quantifying DE or DA without relying on predefined cell clusters [19, 20, 21]. These methods typically embed cells into a low-dimensional space and then consider small neighborhoods in that space to identify “local” DE or DA effects. A remaining caveat of this approach, however, is that it does not account for the uncertainty that embeddings may have (e.g., as inferred with variational autoencoders (VAEs) [22]), which can be substantial [23]. Another line of work uses VAEs to learn the effect of sample covariates on the latent embedding of cells [24, 25, 26]. The primary limitations of this approach are that it assumes the effects they evaluate are constant, meaning they are identical for all cells irrespective of their state, and that they do not account for the uncertainty in estimating these effects.

To address these challenges, we introduce MrVI (Multi-resolution Variational Inference), a probabilistic framework for large-scale (multi-sample) single-cell genomics. For exploratory analysis, MrVI identifies sample groups without requiring *a priori* clustering of the cells. Instead, it allows for different sample groupings to be conferred by different subsets of cells that are detected automatically. For comparative analysis, MrVI enables both DE and DA in an annotation-free manner and at high resolution while accounting for uncertainty and controlling for undesired covariates, such as the experimental batch. The notion at the basis of MrVI is that of counterfactual analysis, which aims to infer what would the gene expression profile of a cell be had it come from a certain sample. This approach provides a principled methodology for estimating the effects of sample-level covariates on gene expression at the level of an individual cell. It relies on a hierarchical deep generative model architecture powered by modern techniques in deep learning, like cross-attention, to model the effects of sample covariates while at the same time providing state-of-the-art performance in the quality of sample integration. On the software side, MrVI leverages the optimization procedures included in scvi-tools [27], allowing it to scale to multi-sample studies with millions of cells.

In the following, we demonstrate that MrVI compares favorably to common approaches for integration, exploratory, and comparative analyses and then showcase its utility in several multi-sample studies. In a PBMC dataset from a COVID-19 study, MrVI identifies a monocyte-specific response to the disease that cannot be directly identified by more naive approaches. In a dataset of drug perturbation screens, MrVI reveals both expected and non-trivial relationships between the assayed compounds. Finally, using MrVI to study a cohort of patients with IBD, we find a previously unappreciated subset of pericytes with strong transcriptional changes in patients with stenosis.

## 2 Results

### 2.1 Multi-resolution variational inference

MrVI is a hierarchical Bayesian model for integrative, exploratory, and comparative analysis of single-cell RNA-sequencing data from multiple samples (e.g., corresponding to human subjects) or experimental conditions (e.g., perturbations in a screen; **Figure 1a**). The model utilizes two levels of hierarchy in order to distinguish between two types of sample-level covariates. The first covariate type captures properties we wish to study in either exploratory or comparative settings - we refer to these as *target covariates*. Typically, an identifier for each sample (e.g., human donor ID or experimental perturbation) is a natural choice for the target covariate to be provided as input to MrVI since it is entirely nested in other sample-level target attributes (e.g., treatment type), thus enabling their analysis. The second covariate type is considered “nuisance”, namely confounders we wish to exclude. In most cases, this will correspond to technical factors such as the sample processing site, library preparation technology, or the study-of-origin in cross-study analyses.

**Figure 1:**
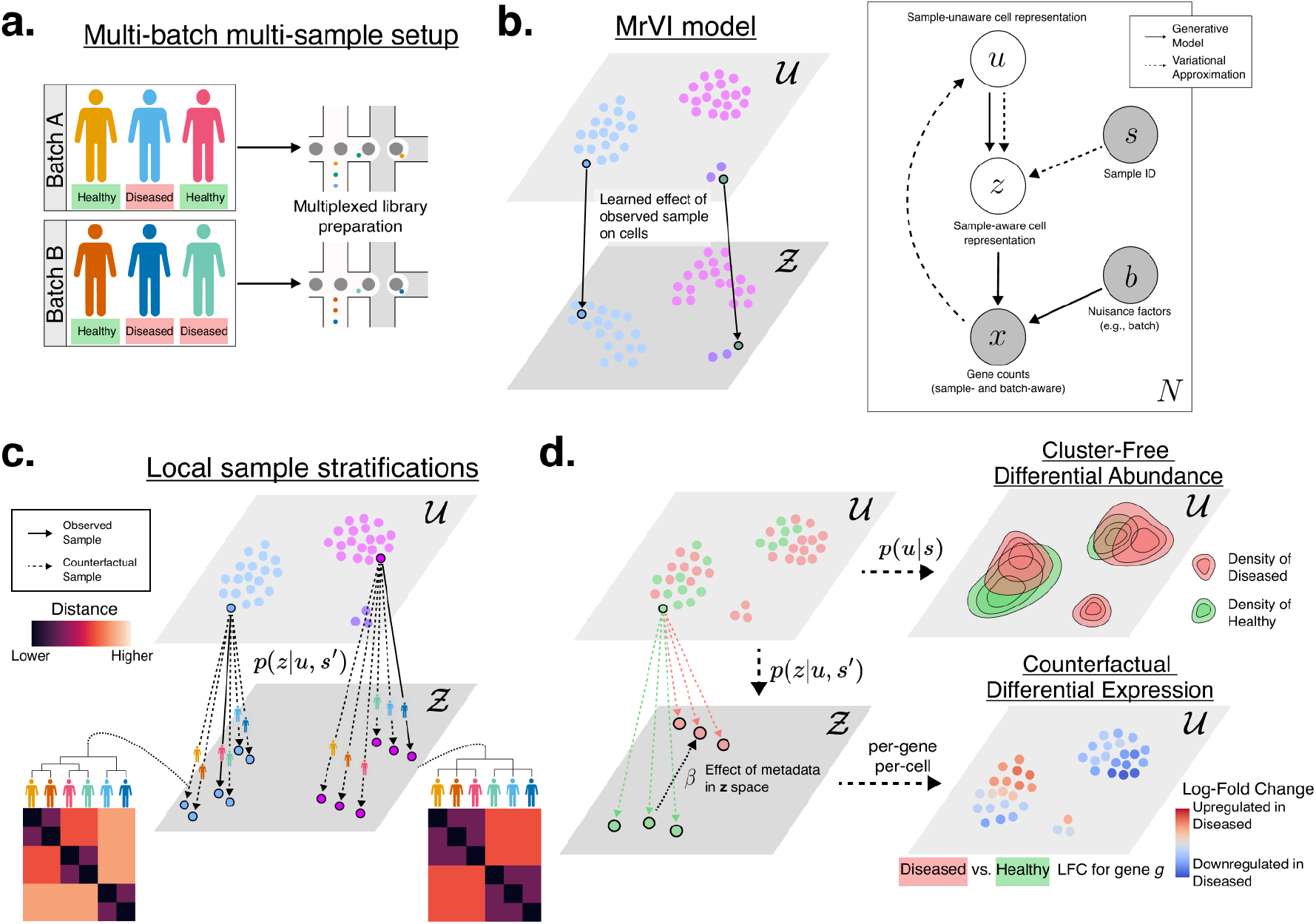
Overview of MrVI. **a**. We consider multi-batch, multi-sample experimental designs. In the canonical case, we gather single-cell measurements from several samples, which are collected across several batches. In this case, the relevant nuisance covariate is the batch. **b**. *(Left)* MrVI model illustration and *(Right)* graphical model plate diagram. A sample-unaware cell representation captures shared type information (colored by cell type in the diagram). From this quantity and the sample-of-origin of the cell, we construct a sample-aware representation of the cell, *z*. Last, we model gene expression as a function of this latent variable and of observed nuisance factors. **c-d**. Use cases of MrVI. MrVI can be used to compute local sample stratifications (*c*), quantify differences in abundance across cell states (*d top right*), and identify sample metadata effects on gene expressions (*d bottom*). Both the sample stratification and differential expression procedures use counterfactual *z* representations to compare local sample effects. The differential abundance procedure involves an approximation of the posterior density for each sample in the *u* latent space.

To formalize this, in MrVI, each cell *n i*s associated with two low-dimensional latent variables, *u*_*n*_ and *z*_*n*_ (**Figure 1b**). The first variable, *u*_*n*_ is designed to capture the variation between cell states while being independent of sample covariates. The second variable, *z*_*n*_, reflects the variation between cell states, in addition to the variation induced by target covariates, while still remaining unaffected by the nuisance covariates. Finally, we model the observed gene expression, *x*_*n*_, as samples from negative binomial distributions whose parameters are predicted by decoding *z*_*n*_ conditioned on nuisance covariates.

Extending our previous work on scVI, MrVI employs a mixture of Gaussians as a prior for *u*_*n*_ instead of a uni-modal Gaussian. We demonstrate that this more versatile prior provides state-of-the-art performance in the integration of large datasets and in facilitating annotations of cell types and states. The distribution of *z*_*n*_ is learned as a function of the respective *u*_*n*_ and the sample ID, *s*_*n*_ (**Methods**). We used neural networks for all mapping functions in the model. The parameters characterizing these functions are learned through maximization of the evidence lower bound (**Methods**) [28].

The trained model can be used to perform both types of analyses - exploratory (de novo grouping of samples) and comparative (evaluating the effects of target covariates) at a single-cell resolution. For exploratory analysis, MrVI computes a sample-by-sample distance matrix, or sample distance matrix in short, for each cell *n b*y evaluating how the sample-of-origin (*s*_*n*_) affects the representation of this cell in *z-*space (**Figure 1c**). To this end, for each cell *n*, we compute *p(z*_*n*_ | *u*_*n*_, *s*′), its hypothetical state had it originated from sample *s*′ ≠ *s*_*n*_. We then define the distance between each pair of samples on cell *n* as the Euclidean distance between their respective hypothetical states. Then, hierarchical clustering can be used over the sample distance matrices for each cell to highlight the target covariates most likely to explain the major axes of sample-level variation. As we show next, this analysis helps capture, in an annotation-free manner, cellular populations that are influenced distinctly by target covariates (e.g., disease or tissue-of-origin).

In comparative analysis, MrVI can identify both DE and DA at a single-cell resolution (**Figure 1d**) using counterfactuals. Consider the case of differential expression between two sets of samples *S*_1_, *S*_2_ as an illustrative example. To evaluate the group-level effects in cell *n*, we evaluate the extent to which the expectation of *p*(*z*_*n*_ | *u*_*n*_, *s*′) depends on whether *s*′ is in *S*_1_ or *S*_2_ using a linear model. We then use the decoder network (i.e., mapping from *z t*o *x)* to detect which genes are affected and evaluate their effect size (fold change). Meanwhile, for local differential abundance, we estimate the posteriors *p*(*u*_*n*_ | *s*′) and compare the aggregate values of samples *s*′ in *S*_*1*_ vs. *S*_*2*_. An in-depth description of MrVI and its post-training analysis procedures are provided in the **Methods** section.

### 2.2 Sample effect characterization in a semi-synthetic experiment

In our first test case, we used a semi-synthetic dataset to evaluate how accurately MrVI captures differences between samples (through exploratory and comparative analysis) when different cell subsets are influenced by different sample-level effects. To generate this, we used a published dataset of 68K peripheral blood mononuclear cells (PBMCs) [29] profiled with 10x, consisting of 3K highly variable genes and five main cell clusters, which we refer to as subsets A-E. We assigned each cell in this dataset to one of 32 synthetic study subjects. These study subjects are characterized by two distinct sample-level covariates. Our strategy for assigning cells to the simulated subjects varied between the different cell subsets so as to simulate different covariate effects. For subset A, the assignment of cells resulted in DE across categories of *covariate 1* in a way that reflects a hierarchical grouping of the samples. For subsets B and C, our cell assignment reflected DA across categories of *covariate 2* (**Figure 2a**). Cells in the remaining subsets were randomly assigned to samples and hence did not contain any DE or DA effects (**Methods**).

**Figure 2:**
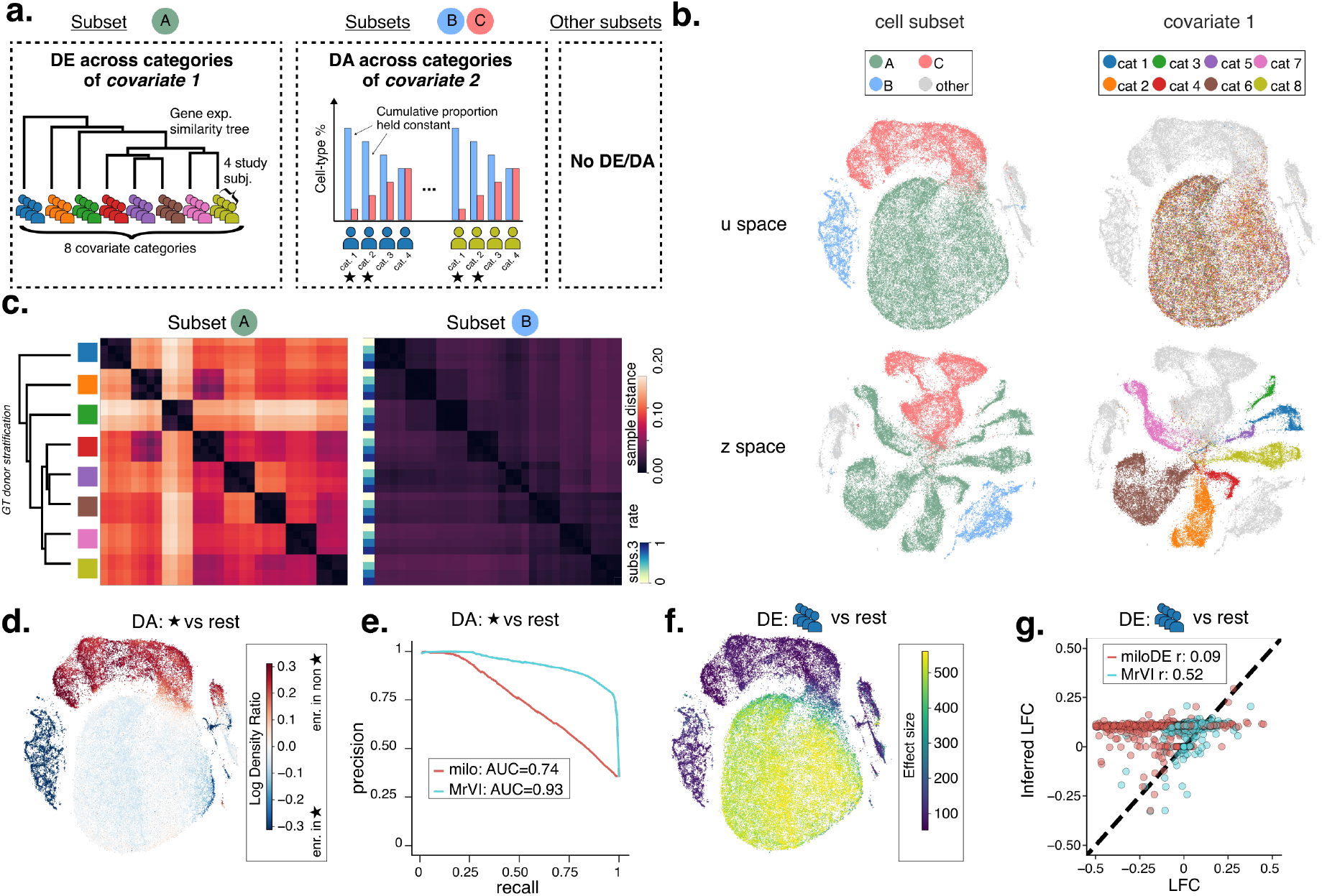
Semi-synthetic experiment. **a**. Experimental design. We generated a semi-synthetic dataset containing 5 subsets of cells and 32 study subjects, with both sample-specific differential expression and differential abundance effects. In cell subset A, cells have differences in gene expression over study subjects based on the value of a study subject covariate, *covariate 1*. In this subset, subjects have differences in gene expression over the categories of *covariate 1* according to a known hierarchy. In two other subsets, B and C, cells have differences in abundance between study subjects based on the value of a second study subject covariate, *covariate 2* (4 categories in total). For each of these categories, cells of subset B (resp. C) were over-sampled (resp. under-sampled) according to fixed rates in each category and in such a way that the sum of cells from B and C remained constant. Stars indicate categories with strong over-sampling/under-sampling. In the rest of the cell subsets, there are no DE or DA effects across study subjects. **b**. Minimum distortion embeddings (MDEs) of MrVI *u* and *z* latent spaces computed on the full dataset, colored by both cell subset assignments (*Left*) and category for *covariate 1* (*Right*). **c**. Cell-subset-specific distance matrices computed by MrVI in the stratified (*Left*) and non-stratified subpopulations (*Right*). **d**. and **e**. Differential abundance analysis using MrVI and Milo. We compared synthetic samples we knew had similar proportions of subsets B and C to those with strong differences in proportions, characterized by stars in **a. d**. MDE of the *u* latent space of MrVI colored by the log density ratios quantifying DA from Equation 5. **e**. Precision-recall curves with areas-under-the-curve (AUC, higher is better) for MrVI and Milo for identifying cells that are differentially abundant for the comparison of samples based on *covariate 2*. We defined a cell belonging to types B or C as a true positive and a true negative otherwise. We used measures of DA strength produced by each respective method (the absolute value of the log density ratio for MrVI; the absolute value of the LFC produced by Milo) as scores and then computed precision and recall at different thresholds. **f**. and **g**. Differential expression analysis using MrVI and miloDE. We compared the differential expression predicted for cells from synthetic samples assigned to one category of *covariate 2* (blue) with the rest of the samples as in **a. f** shows an MDE of the *u* latent space of MrVI colored by an effect size quantifying the overall predicted effect of the sample covariate on gene expression. This effect size corresponds to the squared norm of *β*_*n*_, appearing in Equation (4). **g** compares the log fold-changes inferred by miloDE and MrVI to reference log fold-changes computed using DESeq2. MrVI LFCs, averaged across all cells from the DE cell subset, better correlated to the reference than miloDE. More details about the score used by Milo and MiloDE in e., f., and g. are provided in the **Methods** section.

We applied MrVI, using the simulated subject identifiers as the modeled target covariate *s*_*n*_ and leaving the nuisance covariate *b*_*n*_ empty. The resulting *u s*pace clearly reflected the differences between the cell subsets (**Figure 2b**). In the *z* space, we observed distinct subject-specific effects in cells of subset A, while cells in the remaining clusters were mixed. This result aligned with our expectations, as only subset A contained DE effects. For exploratory analysis, we used the mapping from *u* to *z* to estimate sample distances for each cell (**Figure 2c**). In cell subset A, the sample distance matrix (averaged over all cells of the subset) produced a hierarchical structure similar to the simulated (ground truth) dendrogram. Consistent with our simulation strategy, MrVI estimated much smaller distances between samples when considering the other cell subsets, with no discernible structure. We compared this result to the standard approach for stratifying subjects using differences in cell-type frequencies (**Supplement C**). Specifically, we sub-clustered each subset using cell embedding derived with either PCA or scVI. Then, separately for each subset, we estimated the distances between subjects using the respective sub-cluster proportions. Both standard compositional analyses were less effective in capturing the hierarchy of study subjects in cell subset A (**Figures S1a** and **c**) and were more likely to introduce non-negligible distances in subsets where no differences were expected (**Figure S1b**).

To evaluate MrVI for DA, we partitioned the subjects into two groups according to *covariate 2* (presence or absence of a star in **Figure 2a**). We used the estimated posteriors *p*(*u*|*s*) around each cell to evaluate the extent to which its state was over-represented in one group of study subjects versus the other. The resulting log ratios accurately reflected the DA effects that were simulated in cell subsets B and C (**Figure 2d**). Furthermore, the inferred ratios significantly diverged from zero only in subsets B and C (**Figure S1d**). We compared MrVI to Milo [19], a popular framework for DA. We observed that MrVI more accurately identified the DA effects and associated them with the correct cell subsets (**Figure 2e** and **Supplement C**).

For DE analysis, we compared the subjects in one category of *covariate 1* (blue in **Figure 2a**) to all other subjects. In this comparison, only cell subset A was expected to contain DE effects. We used the estimated posteriors *p*(*z*|*u, s)* around each cell to evaluate the extent to which its gene expression profile depends on the category of its sample-of-origin. The resulting effect sizes were inferred for each latent dimension in *z* using a linear model (see **Methods**; *β*_*n*_ in Equation 4). The squared norm of these effect sizes (aggregating all dimensions of *z)* was used as a measure of the overall effect of covariate 1 on gene expression in each cell.

We observed that these quantities reached much higher values in cells of subset A compared to the remaining subsets (**Figure 2f**). This indicates that MrVI can capture the particular groups of cells exhibiting DE effects. Next, we used the decoder function, *p*(*x*|*z, b*), to evaluate each gene’s respective effect sizes (log-fold changes; LFC) in each cell belonging to subset A. We compared these to effect sizes obtained by a pseudo-bulk DE analysis of subset A (the latter representing an annotation-dependent analysis in the ‘’perfect” scenario where the annotations completely align with the DE signal). We find that the two evaluations of effect sizes were highly correlated, with a substantial improvement over miloDE - a recent method for cluster-free DE analysis (**Figure 2g**).

These results demonstrate that MrVI can identify different sample groupings for different cell subsets without requiring an *a priori* annotation of cell states. Similarly, it is able to accurately retrieve shifts in cell state composition (DA) and gene expression (DE) and identify the respective cellular populations.

### 2.3 Analysis of variation among COVID-19 patients reveals Myeloid-specific stratification into clinical groups

We next employed MrVI to analyze 419k PBMCs obtained from a cohort of COVID-19 patients and healthy controls [7]. We used the sample identifier, corresponding to unique study subjects, as our modeled target covariate *s*. As anticipated, the resulting *u s*pace is not affected by the sample-of-origin, and instead shows marked mixing between study subjects (**Figure 3a**). At the same time, the *u* space clearly stratified the cells into immune subsets in a manner consistent with their annotation in the original study. Considering a standard evaluation of integration performance in this case (using scIB-metrics [30]), we find that the MrVI *u* space embedding outperformed PCA and scVI in terms of mixing the samples while retaining their biological signal [31] (**Figure S2**). To effectively use this dataset, however, we would like our model not to remove the differences between samples but, in this case, to capture the effects of viral infection. While this was not readily achievable with existing integration methods, the two-level structure of MrVI allowed us to derive both a representation that is cell-type-centric (*u*) and one that is affected by the respective sample (*z*). Indeed, the *z* space showed clear sample-specific variation, separating COVID-19-positive patients from the control population inside each cell type (**Figure 3b**).

**Figure 3:**
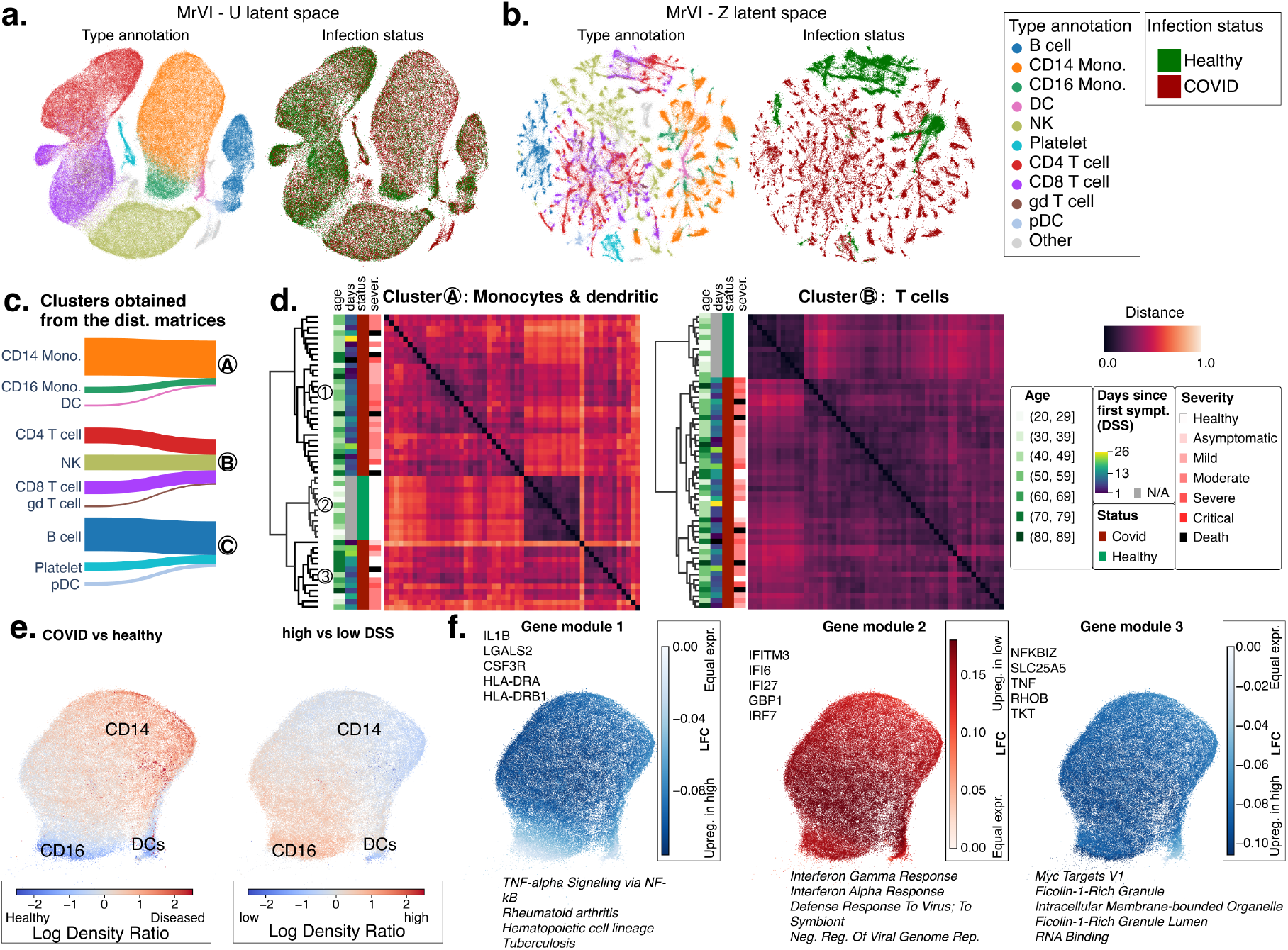
Analysis of a COVID-19 cohort with MrVI. **a**. and **b**. Minimum distortion embeddings (MDEs) of *u* and *z* latent spaces from MrVI, computed on the full dataset and colored by the original cell-type annotations and COVID-19 status. Legends for cell-type annotations and COVID-19 status are shared and displayed on the right of b. **c**. Sankey plot mapping cell-type annotations to clusters obtained by clustering cell-specific distance matrices using the Leiden algorithm. This clustering identified *three* cell subpopulations, Ⓐ, Ⓑ, and Ⓒ. Cluster Ⓐ contained monocytes and dendritic cells, Cluster Ⓑ contained T cells and NK cells, and cluster Ⓒ contained B cells. Cell-type/cluster pairs with less than 1% of the total cells were not displayed. **d**. Sample distance matrices averaged over cells from two of the three subpopulations identified in c. For each matrix, we computed the associated affinity dendrogram between samples obtained via hierarchical clustering and colored each row (sample) according to the patient age, the number of days since first symptoms (DSS), infection status (whether patient or healthy control), and the most severe stage of disease a patient has experienced. **e**. Differential abundance analysis using MrVI log density ratios for the myeloid cells identified as cluster Ⓐ in c. *Left*: Comparison of COVID-19 positive patients to healthy controls. *Right*: Comparison between COVID-19-positive patients with high and low DSS. **f**. Differential expression analysis using MrVI between COVID-19-positive patients with high and low DSS. MrVI identified three DE modules of genes. Each plot shows the activity of the module in the *u* latent space. Displayed are the LFCs averaged over all genes in the module. In these figures, the low and high DSS patients respectively correspond to donor clusters ➀ and ➂ in d.

The two-level representation in MrVI enables an exploratory analysis through two fundamental questions: How do samples in this cohort stratify into groups? And - do they stratify differently when considering different immune populations? To address this, we used the counterfactual embeddings *p(z*_*n*_|*u*_*n*_, *s*′) to estimate a sample distance matrix for each cell *n*. We then clustered the cells according to these values, thus detecting groups of cells that confer similar groupings of the samples and are thus similarly affected by sample-level covariates (**Figure 3c**). Using this analysis, we clustered the cells into three groups, the first containing T cells and NK cells, the second consisting primarily of monocytes along with a smaller population of dendritic cells, and the third containing B cells.

The resulting distance matrices, averaged across all cells in each respective group, provided a clear separation between the patient population and healthy controls, indicating that MrVI can identify clinically relevant groups (**Figure 3d**, **S3**). However, the distance matrix conferred by monocytes and dendritic cells highlighted an additional stratification of the study subjects. In this cell cluster, patients with COVID-19 were further stratified into two groups, corresponding to patient groups ➀ and ➂ in **Figure 3d**. Patient group ➀ was enriched in individuals for whom the number of days since first symptoms (DSS) was low, while individuals ➂ showed longer duration of symptoms (**Figure S4**; Mann-Whitney U test, *p < 0*.*0*5). The association with monocyte activity and the time elapsed since infection has been established [32]. MrVI was able to identify this association without prior knowledge of the DSS or any other information about the human subjects in this study.

To further interpret this data-driven stratification of COVID patients and its association with monocytes, we turned to DA and DE analyses. First, a DA analysis of the myeloid population, comparing the patient population to the healthy controls, showed a marked decrease in non-classical CD16+ monocytes and dendritic cells (**Figure 3e**, **S5a**) in patients. The comparison of the two groups of patients similarly showed a shift toward non-classical monocytes in the group with higher DSS (**Figures 3e**, **S5b**). These results are consistent with independent analysis of COVID-19 patients [32], which reported that CD14+ monocytes are highly pro-inflammatory and contribute to the cytokine release in early COVID, thereby contributing to disease symptoms.

Next, we applied our DE analysis to compare the two patient groups. Using MrVI counterfactuals we estimated the respective effect size (LFC) for each gene in every myeloid cell. We then clustered the genes based on their estimated LFC profiles (see **Methods**). This analysis uncovered three modules, each containing genes with a similar DE pattern implicating different subsets of myeloid cells (**Figures 3f**, **S6**). The first module, upregulated in the group of patients that had higher DSS, was enriched in genes that we also identified in myeloid cells of healthy individuals (compared to patients), again supporting the notion of a return to baseline in myeloid cells with long-standing infection. Specifically, we see a lower *CSF3R* expression in the recently-infected patients, which aligns with less mature monocytes that are released earlier from bone marrow during infection [33]. Similarly, our results indicate that early in the infection, the number of MHC-II-expressing monocytes declines but later returns to normal levels [32]. This accounts for the observed elevation in LGALS2 and HLA-DR2, both of which are linked to MHC-II. The second module, over-expressed in patients with lower DSS, is enriched in interferon-related genes. Specifically, this module includes *GBP1* and *IFITM3*, which are interferon-response genes, and *IFI27*, which was reported as an early predictor of COVID-19 severity [34]. These results agree with a strong interferon signaling during early infection, especially in myeloid cells [32]. The third module, over-expressed by the higher DSS group, contained *TNF* and *NFKBIZ*. It has been demonstrated that TNF release is reduced in acute COVID, whereas *NFKBIZ* is drastically reduced in acute infection [35]. Our analysis suggests that both molecules are markers of acute infection more so than mortality.

Taken together, these results demonstrate that MrVI can simultaneously identify clinically meaningful groupings of study participants as well as the cell subsets that induce them. MrVI also provides the tools to interrogate these stratifications through cluster-free DE and DA analysis. Applied to a cohort of COVID patients, we recovered known compositional changes in the blood myeloid compartment (including a decrease of dendritic cells and non-classical monocytes) as well as gradual gene expression changes in these cells over the course of the disease. MrVI was able to highlight the variation that ensues with DSS as a phenomenon that is more clearly associated with myeloid cells, rather than T or B cells. While the reason for this is currently unclear, monocytes have a relatively shorter half-life time in blood and are replaced by bone marrow-derived cells [36].

### 2.4 MrVI enables grouping and characterization of small molecules in screening assays

To demonstrate the flexibility of the notion of the target covariate in MrVI, we analyzed a chemical perturbation screen with a single cell RNA-seq readout generated with the sci-Plex assay [6]. The sci-RNA-seq3 dataset includes three cell lines and 188 small molecule drugs plus vehicle controls. Each small molecule was delivered at four different doses and the entire study was conducted with two biological replicates. The sci-Plex assay is designed such that each cell receives a single perturbation (or negative control vehicle) that can be identified in addition to its transcriptome. MrVI can serve several fundamental analyses for this type of study, namely integrating all replicates into a shared embedding, stratifying the screened compounds into groups with similar effects, and mapping out these effects at the level of gene expression and the composition of cell states.

To achieve this, we used the concatenation of the drug name and the dose level as the target covariate modeled by MrVI, resulting in 752 “samples” per cell line. Since the study was conducted with 96-well plates, with each plate containing one of the two biological replicates, we chose the plate identifier as our nuisance covariate. As found in the original study, many of the drug-dose combinations had minimal effect on transcription across the assayed cell lines [6]. For this reason, we applied a simple filter, only retaining drugs that had a minimal number of differentially expressed genes with at least one concentration and in at least one cell line (using a standard t-test between cells from each sample against the vehicle; see **Methods**). This resulted in a total of 368 perturbation samples (92 drugs with four concentrations each) that we used for training MrVI (**Methods**). In the following, we focus on the epithelial A549 lung adenocarcinoma line. We provide the results from the other two cell lines in Supplementary Figures S14-S23.

The resulting *u* space (**Figure 4a**) appears as a single cluster with no apparent sub-clusters specific to a given class of drugs, indicating a successful integration that reflects the drug-independent states of the cells. We rather observed that positioning in the *u s*pace carries information about the phase of the cell cycle. With respect to the sample-affected (*z)* space (**Figure 4b**), we observed several sub-clusters of cells originating from distinct classes of drugs. In particular, populations of cells treated with HDAC inhibitors (expected to target epigenetic regulation) and trametinib (a block to MEK-mediated tyrosine kinase signaling) formed clear clusters, highlighting their distinct drug-induced shifts in gene expression. The distinction of HDAC inhibitors is concordant with the original study, in which the authors additionally identified acetyl-CoA deprivation as a common mechanism for this drug class, captured by drug-induced shifts in gene expression. The response to trametinib is also expected to greatly impact the Ras-driven A549 cells since the drug inhibits the downstream MEK pathway [6].

**Figure 4:**
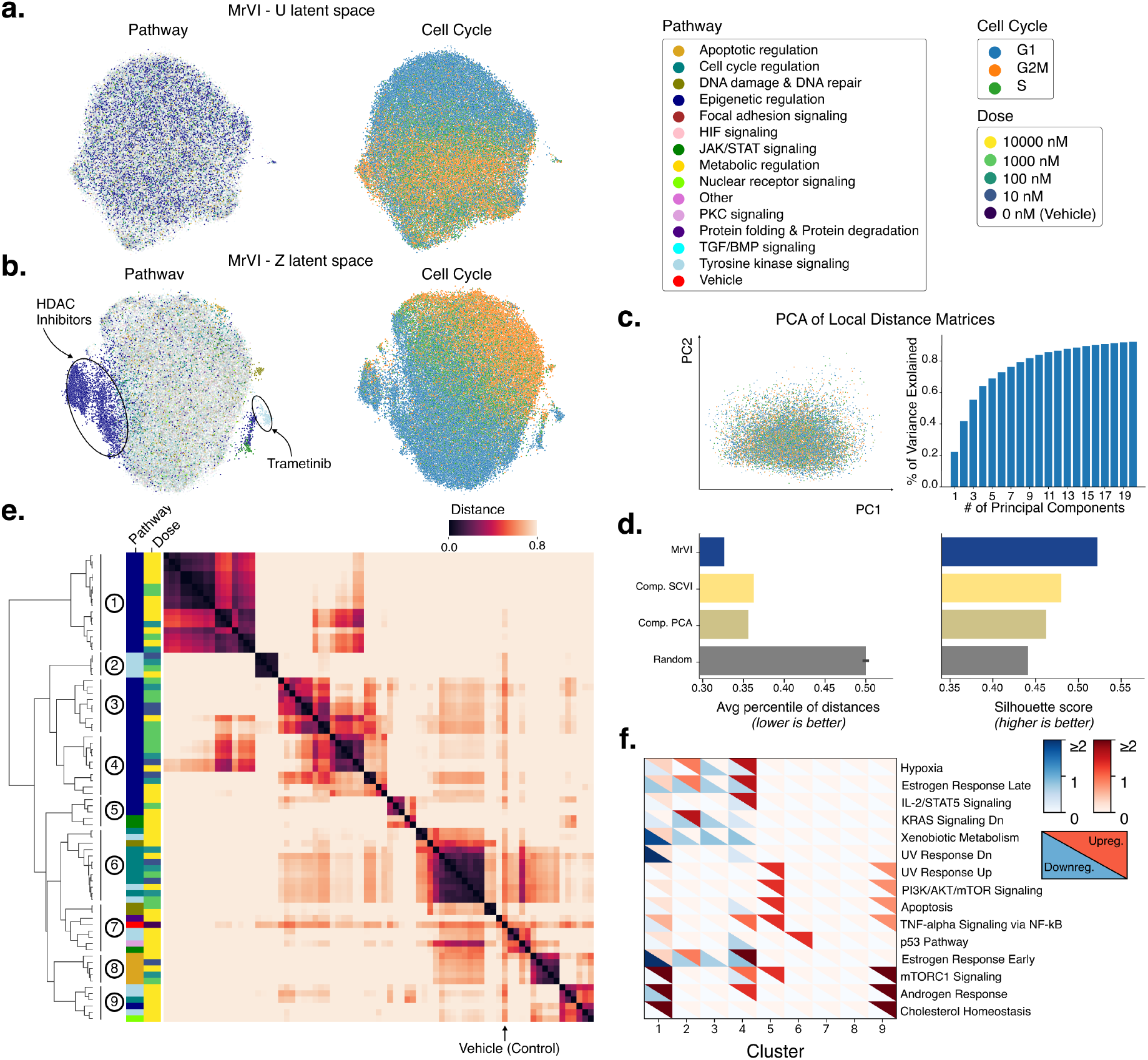
Analysis of the A549 cell line in the sci-Plex experiment. We fit the model over 92 drugs each at four doses that passed our simple DE-gene filter. **a. and b**. MDEs of the *u* and *z* latent spaces from MrVI colored by the target pathway of the drug used to treat each cell (*left*) and the cell cycle stage of each cell (*right*). For the MDEs colored by target pathway, only the top 20 percent of samples based on distance from the vehicle are shown in full opacity. We considered this 2D visualization to be interchangeable with UMAP. **c**. PCA of sample distance matrices. *Left*: scatterplot of all local sample distance matrices projected onto the top two principal components colored by cell cycle stage displays no visual subclusters. *Right*: bar plot of the proportion of variance explained against the number of principal components used. **d**. Barplots comparing MrVI against the benchmark methods for two performance metrics that determine alignment with prior knowledge. *Left*: The average percentile of distances measures how much closer samples with the same drug and different doses are to each other relative to the rest of the distances. We expect the average percentile to be low. *Right*: The silhouette score of sample clusters with similarities inferred from DEG sets in the Connectivity Map dataset. This metric measures whether the clusters are consistent. **e**. Hierarchically clustered sample distance matrix. The rows of the distance matrix are annotated by the pathway and dose of the respective sample (drug-dose combination) and by the clusters inferred from the distance matrix. **f**. Heatmap of Gene Set Enrichment Analysis (GSEA) scores for the Human MSigDB Hallmark gene set collection for differentially expressed genes found for each cluster found in panel e. with respect to the vehicle cells. The upper-right triangle of each tile represents the score for the set of up-regulated DE genes, and the bottom-left triangle represents the score for the set of down-regulated DE genes. For e. and f., the analysis is performed over the top 20 percent of drug-dose combinations (74 out of 368) based on their distance from the vehicle (see **Figure S10** for the full matrix and **Figure S9** for the matrix shown here except with drug-dose labels). Legends for a., b., c., and e. are found in the top right of the figure.

For a more quantitative comparison of the chemical perturbations, we turned to the sample distance matrices estimated by MrVI. We found that the cells in this cell-line-based assay are homogenous in terms of the distance matrices that they induce (**Figure 4c**) with no evident subclusters in PC space. This is in contrast to the other datasets analyzed here that explored primary cells with diverse cell types, featuring distinct distance matrices (**Figures 3**,**5**). Therefore, we operated on one sample distance matrix for the remainder of the analysis, taking the average over all cells.

To test whether the resulting distance matrix captures *a priori* known relationships between the samples, we formulated two performance metrics and compared MrVI to the two standard (composition-based) methods as before (**Figure 4d** and **Supplement D**). First, we used the transcriptomic-based Connectivity Map resource [37], which provides a measure of similarity between drugs, and compared these similarities to the distances we estimated using MrVI for the maximum dose tested (10000 nM). Second, we evaluated the extent to which treatments with the same drug but at different concentrations tend to be more similar to each other than expected by chance. MrVI achieved better performance on both metrics - showing higher concordance with the Connectivity Map stratification and lower distances between treatments with the same compound. Notably, these metrics were also used for fine-tuning the hyperparameters of the MrVI model (here, the dimensions of *u* and *z*). This strategy reflects real-world cases where prior knowledge of the similarity between samples can be utilized for more effective modeling and downstream analysis (**Figure S8**).

To gain additional insight, we analyzed a hierarchical clustering of the sample distance matrix (**Figure 4e**, **S9**). We found that drug-dose combinations that had little to no effect in the A549 context mostly clustered together with the vehicle treatment (**Figure S10**). We observed that disproportionately many of the samples with low effect sizes also have low dosages, capturing expected dose-response relationships. The remaining samples that were most distinct from the vehicle sample were organized into several clusters, each with a different effect on gene expression (**Figures S11**, **S12**). Specifically, clusters ➀, ➂, and ➃ consist mostly of HDAC inhibitors. These three clusters stretch across a wide range of effect sizes that are correlated with the dosages, with cluster ➀ containing the samples with the highest dosage levels and the largest effects and cluster ➂ consisting of samples with lower dosages and weaker effects (**Figure S11**). These groupings highlight the ability of MrVI to uncover dose-dependent effects on gene expression apparent across multiple drugs in the HDAC inhibitor class. Clusters ➁ and ➇ corresponded to all doses of trametinib and YM155, respectively. In these cases, MrVI therefore suggests that the effects of these drugs were less dependent on the dose, at least when considering the range of concentrations tested.

While the clusters aligned with the expectation that drugs labeled to target the same pathway or different doses of the same drug should have similar effects, MrVI also uncovered relationships between drugs based on their effects on transcription that are supported by recent literature and the original sci-Plex study. For instance, cluster ➅ includes rigosertib, labeled as a tyrosine kinase inhibitor but found to directly affect microtubule function [38], as well as epothilone A and patupilone, two drugs that interfere with the microtubule function. Moreover, MrVI revealed non-trivial similarities that were not captured in the original study. In cluster ➄, two JAK2 inhibitors, fedratinib and TG101209, were grouped with JQ1, a drug labeled as a BRD inhibitor. Interestingly, recent work has shown that JQ1 inhibits the JAK-STAT signaling pathway in addition to being a BRD inhibitor, which supports the plausibility of this grouping [39].

Finally, we investigated the clusters by performing Gene Set Enrichment Analysis (GSEA;[40]) on the DE gene sets identified by MrVI (comparing each cluster of samples to the vehicle controls; **Figure 4f** and **Methods**). As a reference, we used the hallmark collection of MSigDB [41] that records sets of genes that contribute to major cellular processes. This analysis shed additional light on the effects of each cluster of drugs. Specifically, we found that clusters ➀, ➂, and ➃ associate most strongly with a down-regulation in metabolic pathways, agreeing with the effect of HDAC inhibitors on carbon metabolism [6]. Furthermore, Cluster ➅ was enriched in the p53 pathway, consistent with the categorization of its respective drugs as cell cycle regulators. Similarly, the effects of Cluster ➁ were enriched in genes downstream of KRAS signaling, consistent with its categorization as targeting tyrosine kinases. We provide a heatmap of the LFCs for the top DE genes across all clusters in the supplement (**Figure S13**). Based on this analysis, we highlight that MrVI not only provides an interpretable grouping of clusters but additionally helps highlight the genes underlying this grouping.

For the other two cell lines used in the sci-Plex experiment, the results of MrVI were consistent with known biology (**Figures S16-S23**). For instance, the hormone-receptor-positive breast cancer cell line, MCF-7, was the only cell line to exhibit strong effects in response to the hormone therapies, fulvestrant and toremifene citrate, as evident in both the *z s*pace and the sample distance matrix (**Figure S14**). In particular, fulvestrant, which had a strong effect at the highest two dose levels on MCF-7 [42], had little effect on other cell lines and did not appear in the top 20% of drug-dose combinations for the other two cell lines based on distance from the vehicle. The GSEA results also showed strong down-regulation of estrogen-response-related genes in response to many of the drugs. Similarly, in the Bcr-Abl positive cell line K562 [43], we observe a cluster of Bcr-Abl tyrosine kinase inhibitors (bosutinib, dasatinib, nilotinib) with a significant effect absent from the other cell lines (**Figure S15**).

Overall, MrVI recapitulated both expected and novel aspects of the effects of the screened drugs and the relationship between them. It did so through an end-to-end solution that coupled integration across experiments, estimation of the effects of individual molecules, stratification of the affecting molecules into biologically meaningful groups, and characterization of the effects of each group. MrVI uncovered non-trivial relationships between drug dosage and effect on cell lines, highlighting dosage-independent and -dependent effects. More broadly, this analysis demonstrates the utility of MrVI in powering screening studies (e.g., genetic or chemical) that leverage single-cell readouts for studying large numbers of perturbations.

### 2.5 Characterizing the role of stromal cells in stenosis in Crohn’s disease

To provide another example of the applicability of MrVI to human cohorts, we utilized a recent study of Crohn’s disease, conducted in 46 patients and 25 controls using single-cell RNA-sequencing [44]. In addition to gene expression, this dataset comes with metadata describing the anatomical location of sampling (in terms of tissue: colon or ileum, and in terms of the tissue layer: lamina propria or epithelial), the method of extraction (surgical or biopsy), and sample preparation detail (10X chemistry). It also includes information on the respective study participant, such as disease state and the presence of stenosis in the patient’s history (i.e., narrowing of the intestines due to inflammation). We used these metadata to evaluate the ability of MrVI to recognize meaningful patient sub-groups and to highlight cell populations that are affected by stenosis.

We trained MrVI to integrate all the samples in this dataset, using the sample identifier as the modeled target covariate and the combination of library preparation protocol and tissue layers (lamina propria and mucosa) as the nuisance covariate. Comparing the resulting *u* space to the embedding obtained with scVI, we found that the default settings of MrVI yielded better mixing between the study subjects but had slightly lower performance in terms of distinguishing between cell states (using cell annotations assigned in the original study; **Figure S24**). Indeed, integration is challenging in this dataset due to significant differences in cell type composition in the colon and in the ileum, as well as between the mucosal and lamina propria layers. To increase the extent to which the *u* representation captures variation between cell states, we developed a variant of MrVI that makes use of cell type labels. Specifically, we introduce a cell-type-specific bias term for the mixture weight of the mixture of Gaussians prior used by MrVI, thus encouraging similar embedding for cells of the same state (**Methods**). Compared to scVI, this added consideration of cell type labels leads to an overall improved performance, considering mixing between samples and preserving cell type information (**Figure S24**).

We next used MrVI to explore how the different samples stratify and how these strata change between cell types. Using a coarse definition of cell types (**Figure 5a** and **Methods**), we again find that different types are associated with different patient groupings. For instance, considering a subset of immature Enterocytes (referred to herein as Enterocytes-Stem), the sample distance matrix clustered solely by their tissue-of-origin (colon or ileum) (**Figure S25**). We applied the DE function of MrVI on the Enterocytes-Stem subset to investigate the differences between these clusters of samples. In line with their biological functions, we find higher expression of *AQP8, CA2* and *CA8* in the colon, which all encode proteins that absorb water, and a higher expression of *FABP6* and *FABP2* in the ileum, which encode proteins that absorb fatty acids. Considering a population of mature Enterocytes provides a slightly different view, highlighting a specific cluster of eight patients that are distinguished from all other patients and controls. Using MrVI DE to compare mature Enterocytes in this cluster, which mainly contained colon samples, to other colon samples, we detected an up-regulation of mucins (*MUC1, MUC2, MUC12*), which is a well-described pattern in Crohn’s disease [45], and an up-regulation of *CXCL1* and *CXCL3*, which encode chemokines that attract neutrophils [46]. The expression of these chemokines was associated with stimulation of epithelial cells with *IL22* and *IL17A* that are key cytokines in Crohn’s disease. Furthermore, neutrophil infiltration is a key feature of inflamed gut regions [47]. We find here that, depending on the cell subset under consideration, the exploratory analysis of MrVI can reflect known differences between the tissues sampled (here ileum vs. colon) as well as reveal differences in a subpopulation of diseased individuals (non-inflamed vs. inflamed).

**Figure 5:**
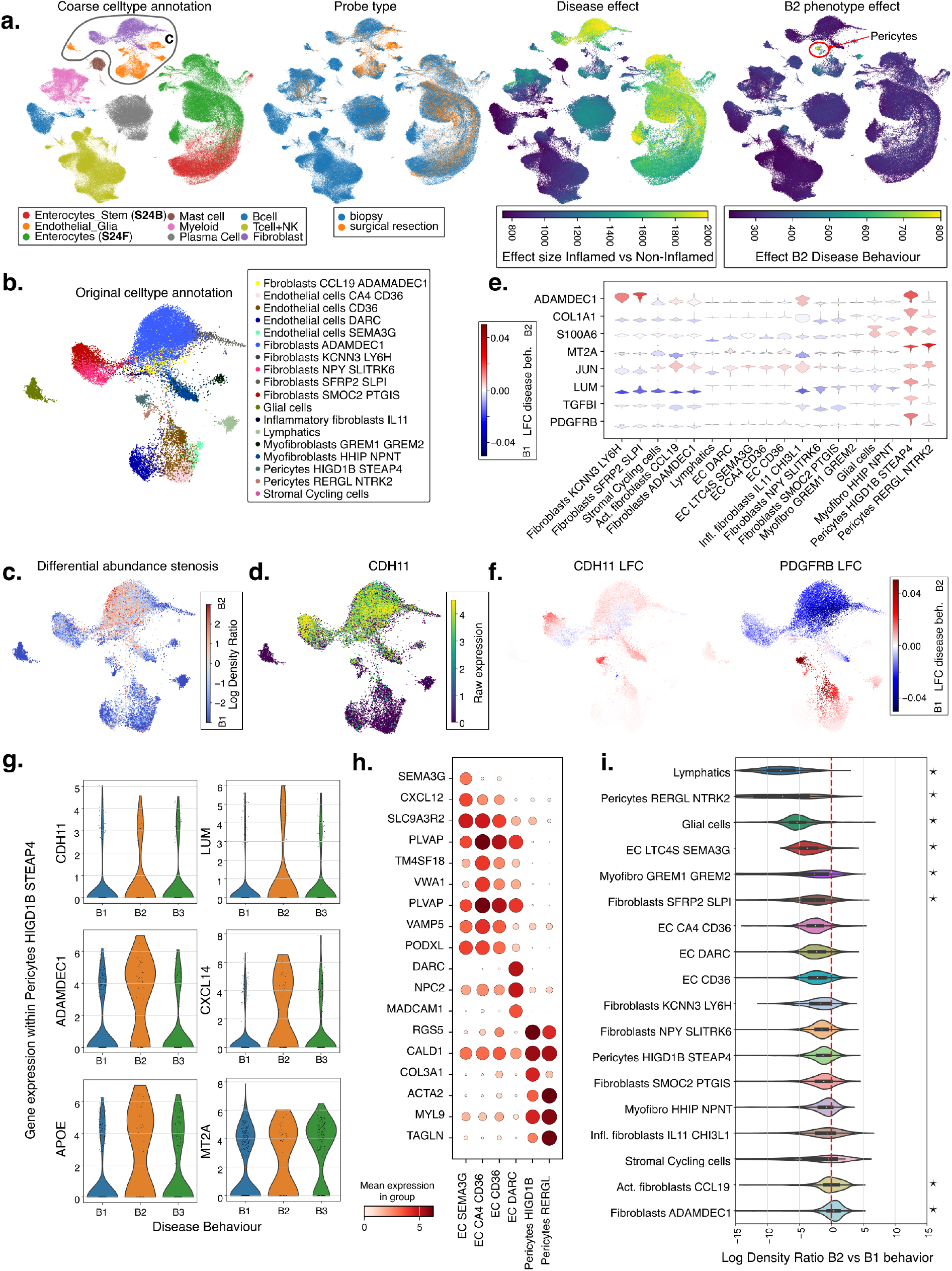
Characterization of stenosis in Crohn’s disease. **a**. UMAP embedding of *u* latent space colored by different sample-level covariates. From left to right: coarse labels identified by us to stratify cell types for unguided analysis (circled is the subset analyzed in **b-e**); tissue collection method highlights a bias for stromal cells in surgical specimens; the inferred effect size of inflamed vs. non-inflamed and the inferred effect size of B2 disease behavior (highlighted are the small subpopulations of pericytes with the strongest effect (**Figure S26**). For these last two plots, this effect size corresponds to the squared norm of *β* from Equation (4); it characterizes the overall effect of the covariate on gene expression. **b-e**. Analysis of the cell population circled in a, using the same UMAP embeddings as a. **b**. Original cell-type labels provided in the original study. Displayed are fine annotations for stromal cells and coarse labels for the other cells. **c**. DA analysis of stromal cells based on MrVI comparing the B2 disease phenotype against the B1 phenotype (red denotes higher in B2 disease behavior). Displayed is the log density ratio between B2 and B1 disease behavior. **d**. Raw expression of *CDH11* inside all stromal cells. Values are library-size normalized and log1p transformed. Cells are sorted for display based on higher expression. **e**. Inferred LFCs from MrVI for the comparison of B2 and B1 disease behavior based on multivariate analysis in MrVI while correcting for inflammation status, sex, tissue location, and chemistry in the different cell types. The violin plots display the distribution of estimated LFCs per cell type. We relied on hierarchical clustering to determine the order of cell types displayed in the violin plots. **f**. UMAP of two genes highlighting intra-cell-type DE variation. Cells are colored based on the genes’ LFC for the comparison of B1 and B2 classifications. **g**. Violin plots illustrating changes in normalized gene expression between inflamed and non-inflamed biopsies from patients categorized as B1, B2, and B3 disease behavior after subsetting to *Pericytes HIGD1B STEAP4*. Raw gene expression values are normalized by library size and log1p-transformed. **h**. Dotplots displaying top 3 marker genes for the different endothelial cell (EC) subclusters. Lymphatic endothelial cells were excluded from this analysis. **i**. Differential abundance from **c** displayed as violin plots to demonstrate changes in abundance across cell types. Displayed is the log density ratio between samples from B2 and B1 disease behavior. We report the significance of the hypothesis that the difference of log density ratios between a given cell type and all other cell types is above 1 in absolute value (see **Methods**).

Next, we explored the use of MrVI for comparative analysis with respect to a known covariate. We consider the distinction between the two most common complications of Crohn’s disease: stenosis (Vienna classification B2) and fistula or abscesses (penetrating; Vienna classification B3). These are evident in 11 and 7 of the subjects, respectively, while the remaining are healthy controls or patients without either complication (Vienna classification B1). Finding reliable biomarkers that can distinguish between patients experiencing the two types of complications is critical, as they may require different treatment strategies. Using our multivariate DE procedure, we evaluated the extent to which each individual cell is impacted by the presence of stenosis as well as by the inflammation status (inflamed vs. non-inflamed) while accounting for the effects of nuisance covariates such as biological sex and tissue location (**Methods**). We excluded surgical samples from this analysis due to the marked differences in cell type composition compared to biopsies (which are the source of most cells in the dataset; **Figure 5a** and **Methods**). We find that inflammation status had a marked effect on several cell lineages, with a strong effect on the stromal compartment. As expected, the presence of B2 disease had a more mild effect, mostly restricted to a few stromal subsets, with its highest impact in a small subset of pericytes (**Figures 5a, S26** and **S27b**). Therefore, we focus the remainder of our analysis on stromal populations consisting of fibroblasts, pericytes, glia cells, and endothelial cells (**Figures 5b**).

We first compared B2 vs. B1 samples using DA analysis while controlling for inflammation status and other covariates (**Methods**). We find a decrease in the abundance of several endothelial populations in B2 samples (e.g., lymphatic and *LTC4S*+ endothelial cells) and an increase in fibroblast populations (e.g., *ADAMDEC1*+; **Figures 5b, c, h**). This result accords with the prevalence of microvascular rarefaction (i.e., loss of endothelial cells) in tissue fibrosis [48]. Using the same settings for DE, we find that in patients classified as B2, a subset of *HIGD1B+STEAP4+* pericytes up-regulated *CDH11*, a biomarker of stenosis [49], as well as classical markers of tissue fibrosis (*ADAMDEC1* and *COL1A1*) and activation (*S100A6, MT2A*, and *JUN*) (**Figures 5d, e, f** and **S27d**). While these genes are up-regulated in B2 disease by other stromal subsets, we find several genes that are affected uniquely in cells of the *HIGD1B+STEAP4+* pericyte subset. This includes *LUM*, which was shown to be up-regulated after fibroblast stimulation in lung fibrosis and promotes fibrocyte differentiation [50] as well as *PDGFRB* and *TGFBI*, which are strongly up-regulated in lung fibrosis and whose pharmacological inhibition reduces fibrosis [20, 51]. Furthermore, *LUM* and *TGFBI* were reported to be up-regulated in the chronic phase of a mouse model of colitis [52], which is associated with marked intestinal fibrosis. Notably, while MrVI bases its DE estimations on its generative model, targeted analysis of the same molecules using the raw data shows consistent results (**Figure 5g**). Together, this analysis demonstrates the potential of MrVI for delineating a population of cells that is associated with a disease phenotype and may facilitate a more nuanced discovery of markers for diagnosis and treatment.

MrVI additionally predicts marked up-regulation of *PDGFRB* by *CD36+* endothelial cells in B2 samples(**Figure 5e, f**). This is unexpected as *PDGFRB* is a common marker of pericytes and is not normally expressed by endothelial cells. We further characterized gene expression in the *CD36+* subset and found co-expression of endothelial markers (like *PLVAP, VAMP5, VWA1*) and pericyte markers (like *NOTCH3, RGS5* and *MYL9*) (**Figures 5h** and **S27d**). In addition, we find up-regulation of markers of tissue fibrosis such as *COL1A1* and *TGFBI* in this subset (**Figure S27e**). Therefore, these results highlight a cell population with a mixed phenotype between the endothelial and pericyte lineages, which up-regulates markers of tissue fibrosis in B2 samples. This hints towards endothelial-to-mesenchymal transition in IBD and suggests the presence of a pericyte-like state in the gut endothelium of B2 disease. Endothelial-to-mesenchymal transition has been described for human IBD [53]. However, this phenomenon has not been explored in the original study of this cohort, nor, to date, studied with single-cell genomics [53].

In summary, MrVI uncovered previously unappreciated changes in cellular and molecular composition in patients with stenosis. It identified cells with strong changes in gene expression in stenosis, detected several genes associated with tissue fibrosis, and suggested endothelial-to-mesenchymal transition in stenosis.

## 3 Discussion

We introduced MrVI a comprehensive solution for large-scale (multi-sample) single-cell RNA-seq studies. MrVI provides a unified probabilistic framework for several fundamental tasks, namely integration of samples from different sources, sample stratification, and analysis of the effects of sample covariates at both the cell-subset and gene levels. Based on a hierarchical latent variable model and counterfactual predictions, MrVI is capable of addressing these tasks while accounting for nuisance sources of variation and estimating uncertainty without the need for *a priori* annotations of cells into types. The latter point is of particular importance due to the difficulty of defining cluster boundaries and their resolution, which can vary substantially across single-cell studies. For instance, [10] categorized the human brain into 17 cell types for studying autism while [54] identified 3313 clusters to characterize cellular heterogeneity in the same tissue. Both strategies proved useful for their respective study, and it is therefore not generally clear which resolution is appropriate and to which type of analysis. Instead of relying on a fixed strategy for clustering of the cells, MrVI facilitates a “bottom-up” approach that divides the cells into groups in a manner that reflects the task at hand. Specifically, by estimating sample distance matrices around each cell, MrVI allows for the aggregation of cells into subsets that confer similar groupings of samples. Similarly, estimating DE or DA effects in every cell allows for the aggregation of genes or cells in a way that reflects a coherent response to the covariate of interest. For ease of interpretation, these aggregations can also make use of *a priori* annotations of cells into subsets (by averaging the cell-wise DE or DA effects), as long as the cells within a subset are consistently affected (which was the case in many of our analyses).

We demonstrated the capability of MrVI to perform these fundamental analyses in a few case studies. Considering a COVID-19 cohort, MrVI identified clinically relevant patient groupings and highlighted a subsets of myeloid cells in which these groupings are manifested. Post-hoc analysis of the resulting patient strata further revealed a marked agreement with the elapsed time since infection - information that was not available to the algorithm. Importantly, this patient grouping did not perfectly mirror the infection timelines. For instance, some subjects with higher DSS exhibited molecular and cellular characteristics akin to those in the recently affected group. This observation does not undermine the validity of our approach; it rather underscores the potential of MrVI to produce data-driven sample strata that may not be trivially obtained from the recorded metadata alone and instead may lead to different diagnoses or identification of new disease sub-types [12]. MrVI is particularly relevant for studies in which samples are collected from numerous individuals, possibly collected across different anatomical locations or experimental protocols. We demonstrated this using a Crohn’s disease study, where it effectively integrated samples from diverse tissue locations and highlighted changes associated with stenosis. MrVI’s utility is not restricted to clinical studies or to other comparisons that are at the level of an experimental sample. Instead, it can be applied to any discrete cell-level meta-data by designating it as the target covariate. We demonstrated this using a perturbation screen with the sci-Plex assay in which each cell is associated with a particular perturbagen, facilitating *de novo* identification of compound groups and characterization of their effects.

When considering patient cohorts, we applied MrVI using the sample identifier as our modeled target covariate, *s*. This design enabled us to study the effects of any sample-level property (which is trivially nested by the sample identifier). Future extensions of MrVI involve adapting the model to account for multiple sample-level target and/or nuisance covariates. For instance, conditioning on sample-level covariates with strong and unambiguous effects (e.g., sex, disease status) could help uncover more subtle effects attributed to other target covariates. Additionally, covariates could help improve integration in the *u s*pace, especially when specific cell states are not shared across all samples. Allowing MrVI to condition on continuous covariates (e.g., drug dosage, time) would also pave the way for inferring cell-state transitions and intermediate states.

While this work primarily focused on scRNA-seq data, a natural extension of MrVI is to handle information from other measurement modalities, both separately and in parallel to RNA expression. Such an extension could pinpoint, for instance, different patient strata (and their inducing cell subsets) when considering chromatin properties vs. RNA [55]. As another extension, adapting existing transfer learning protocols to MrVI [56] could enable the analysis of smaller datasets. In the same way that transfer learning leverages annotated cell atlases to label query datasets, transfer learning for MrVI could provide a way to harmonize samples across studies, especially when essential sample metadata are missing or inaccurate. For example, a MrVI model pre-trained on a large cell atlas with rich sample-level metadata (e.g., age, sex, inflammation status) can be leveraged to provide insight into unrecorded properties of new samples based on how they stratify relatively to the reference samples.

MrVI is implemented using state-of-the-art software tools for deep probabilistic modeling and can thus scale to multi-sample studies with millions of cells. Beyond that, the expected increase in scale and complexity of single-cell omics raise new challenges and opportunities for which MrVI can provide a powerful framework for analysis and a solid foundation for further developments.

## Acknowledgments and Disclosure of Funding

We would like to thank Lingting Shi, Florian Ingelfinger, Fadi Sheban, Nathan Levy, Ross Giglio, Nicholas Hou, Kevin Hoffer-Hawlik, Sopho Kevlishvili, and Avital Steinberg for being the first to try the MrVI Python package and providing valuable feedback that greatly improved our work.

This work was supported by a Chan-Zuckerberg Initiative Seed Networks for the Human Cell Atlas grant (CZF2019-002452) and NIAID Grant R01 AI169075 to N.Y. J.H. was supported by grant number 2022-253560 from the Chan Zuckerberg Initiative DAF, an advised fund of Silicon Valley Community Foundation. A.G. is currently an employee of Google DeepMind. Google DeepMind has not directed any aspect of this study nor exerts any commercial rights over the results.

## Methods

### A The MrVI model

#### A.1 Generative model overview

We consider two-stage scRNA-seq experimental designs in which cells are collected from multiple samples (**Figure 1a**). Each sample is associated with target covariates (e.g., treated vs. untreated, donor or specimen age and sex) or nuisance covariates (e.g., the sample collection site or the study ID in cross-study analyses).

Typically, multiple target covariates can induce variation in expression across samples, but it is unknown which of these may affect cells and by what mechanism. For instance, in drug response studies, both the type of administered drug and its dosage are crucial to assessing drug impact on cell states, but the nature of the interaction between these two factors may not be known. In disease studies, cases may induce specific shifts in gene expression in specific donor subpopulations that may not be fully encoded in the available metadata.

Instead of attempting to model the effects of these covariates directly, we adopt an approach that initially requires only knowledge of sample IDs *s* ∈ {1, …, *S*} and the nuisance covariates as *b* ∈ {1, …, *B*}. This strategy allows us, at a later stage, to highlight which target covariates drive sample variations of interest. The resulting gene expression profiles are denoted as {*x*_*1*_, …, *x*_*N*_}, where *x*_*n*_ ∈ ℕ^*G*^ is the vector of RNA transcript counts for cell *n* over the *G* observed genes. For any cell *n*, we let *s*_*n*_ identify the sample ID (e.g., the donor from which cell *n* originates) and *b*_*n*_, the nuisance covariate.

In the case of multiple nuisance covariates, we recommend using the covariate with the coarsest resolution that is still nested within any covariates expected to confound the analysis. This may require concatenating multiple nuisance covariates (i.e., the study ID concatenated with the batch ID used in each study as *b*_*n*_).

##### Isolating sample-specific effects on cell states with *MrVI*

The generative model of MrVI writes as

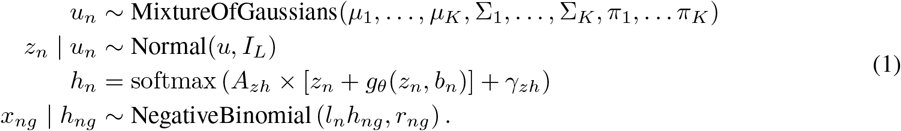

Here, *u*_*n*_ and *z*_*n*_ are the latent (unobserved) representations of cell *n*, both of dimension *L. A*_*zh*_ is a matrix of dimension *G × L* and *γ*_*zh*_ is a bias vector of dimension *G. g*_*θ*_ is a multi-head attention layer, with parameters *θ*. The size factor *l*_*n*_ is fixed as the total sum of counts of cell *n*, and *r*_*ng*_ ≥ 0 denotes the inverse dispersion of the distribution. *µ*_*i*_, *Σ*_*i*_, *π*_*i*_ respectively denote the mean, covariance matrix, and weight of component *i* ≤ *K*. More details about this prior are provided in Section B. All these parameters, other than *l*_*n*_, are learned during training.

We now unpack some key aspects of the model. *u*_*n*_ captures broad variations assumed to characterize cell types and more granular cell states but is independent of both target and nuisance covariates. As such, *u*_*n*_ harmonizes cells from all samples into a shared latent space. We assume a mixture of Gaussians (MoG) prior or an unimodal Gaussian prior (*K =* 1) on *u*_*n*_, depending on the application and available prior knowledge about cell-state variation. When we *a priori* expect cells to belong to one of several groups, a MoG prior may be more appropriate than an unimodal Gaussian prior to avoid posterior collapse and prior overregularization, two issues with variational inference reported in the field [57, 58]. When reliable cell-type annotations are available, MrVI can also rely on a prior that is weakly informed about cell-type annotations. More details about the prior of *u*_*n*_ are given in Section B.

*z*_*n*_ is an augmented representation of the cell, that is aware of sample effects but is independent of other nuisance covariates. This latent variable is constrained to be close to *u*_*n*_ by the isotropic Gaussian prior centered on *u*_*n*_.

As *z*_*n*_ is expected to capture more variability within cells than *u*_*n*_, allowing it to lie in a higher-dimensional space is natural. Additionally, a low-dimensional bottleneck on *u*_*n*_ may improve sample harmonization. In such a case, we allow *z*_*n*_ to take a higher dimension than *u*_*n*_ by modeling *z*_*n*_ ∼ Normal(*A*_*uz*_*u*_*n*_ + *γ*_*uz*_, *1*), where *A*_*uz*_ is a learned matrix of dimension *L*_*z*_ *× L a*nd *γ*_*uz*_ is a bias vector of dimension *L*_*z*_, where *L*_*z*_ is the dimension of *z*_*n*_. Without loss of generality, the remainder of the manuscript focuses on the case where *z*_*n*_ and *u*_*n*_ have the same dimension, *L*.

##### Modeling gene expression under technical effects

MrVI models the normalized expression of gene *g*, denoted *h*_*ng*_, as a function of both *z*_*n*_ and the nuisance covariate. This relationship is parameterized with multi-head attention (*g*_*θ*_ above) to capture nonlinear nuisance-covariate-specific effects on gene expression. More information regarding this parameterization is given in Section B. Lastly, we model the observed transcript counts with negative binomial distributions and account for the technical effects of the sequencing depth using the same approach as scVI [22].

#### A.2 Variational approximation and training procedure

The generative model described by Equation (1) can be used to generate synthetic data; it does not directly inform us on the posterior distribution of the latent variables *u*_*n*_ and *z*_*n*_ given observed gene expressions and sample ID of a given cell *n r*equired for analysis. Since posteriors *p*_*θ*_(*u*_*n*_, *z*_*n*_ | *x*_*n*_, *s*_*n*_) are intractable, we rely on variational inference to learn an approximation *q*_*ϕ*_(*u*_*n*_, *z*_*n*_ | *x*_*n*_, *s*_*n*_) to the posterior, where *ϕ d*enote all parameters used to construct the variational approximation. The variational distributions we consider factorize as *q*_*ϕ*_(*u*_*n*_, *z*_*n*_ | *x*_*n*_, *s*_*n*_) = *q*_*ϕ*_(*u*_*n*_ | *x*_*n*_)*q*_*ϕ*_(*z*_*n*_ | *u*_*n*_, *s*_*n*_). We now describe these in more detail.

##### Modeling *q*_*ϕ*_(*u*_*n*_ | *x*_*n*_)

We model *q*_*ϕ*_(*u*_*n*_ | *x*_*n*_) as a Gaussian distribution whose mean and covariance (assumed diagonal) are outputs of multi-layer perceptrons (MLPs) taking *x*_*n*_ as inputs.

##### Modeling *q*_*ϕ*_(*z*_*n*_ | *u*_*n*_, *s*_*n*_)

The variational approximation to *z*_*n*_ relies on a multi-head attention mechanism [59] taking both *u*_*n*_ and *s*_*n*_ as inputs. We have empirically observed this multi-head attention mechanism to better capture localized sample effects than MLPs, as the latter tended to learn global effects across all cell states even when this effect was localized to a specific subset of the cells. Additionally, we found no practical benefits in modeling the uncertainty around *z*_*n*_. Instead, we used a point mass approximation to the posterior of *z*_*n*_ given *u*_*n*_ and *s*_*n*_. Overall, given *u*_*n*_ and *s*_*n*_, *z*_*n*_ can be obtained as

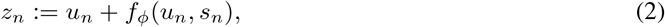

where, here, *f*_*ϕ*_ is the multi-head attention mechanism (the same as for *g*_*θ*_), described more at length in Section B.

With these two components, a straightforward approach to get posterior latent *u*_*n*_ and *z*_*n*_ for a given cell consists of sampling *u*_*n*_ from *q*_*ϕ*_(*u*_*n*_ | *x*_*n*_), then computing *z*_*n*_ as described in Equation (2).

##### Training procedure

We optimize the evidence lower bound (ELBO), which we maximize over the generative model parameters *θ a*nd variational parameters *ϕ u*sing mini-batch stochastic gradient descent methods [60, 61]. In this problem, the ELBO writes as

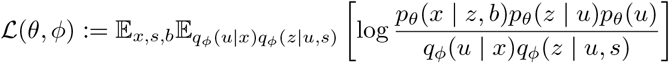

#### A.3 Exploratory analysis of sample effects on cell states

A common scenario in large-scale studies is that sample-level covariates are incomplete, noisy, or inconsistent across datasets. It may also be the case that the most relevant sample characteristics affecting gene expression are unobserved. Here, MrVI can identify the most relevant sources of heterogeneity between samples without assuming access to relevant target covariates (**Figure 1c**); the most salient axes of variation across samples can then be related back to observed target covariates. This type of analysis relies on cell-state counterfactuals, which are used to quantify sample distances at the cellular level.

##### Predicting counterfactual cell states

After model fitting, MrVI can be used to predict the effect of a given sample on any cell. We aim to predict the counterfactual state of the cell, that is, its state had it been collected from another sample. We achieve this by substituting the sample-of-interest *s*′ ≠ *s*_*n*_ for the true sample-of-origin *s*_*n*_ in Equation (2) to obtain

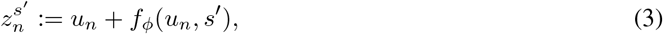

where *u*_*n*_ is the inferred cell state for cell *n o*btained via variational inference. 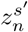 captures the counterfactual state of cell *n h*ad it been collected from sample *s*′.

##### Estimating sample distances at the cellular level

MrVI allows for unsupervised sample stratification by comparing distances between counterfactuals from Equation 3. In fact, we can assess the differences between samples *s*_*a*_ and *s*_*b*_ on a cell *n b*y computing the distance between their respective counterfactual cell states 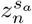 and 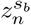. In particular, low distances between counterfactuals indicate that the two samples have similar effects on the cell according to the model.

Based on this observation, we summarize the sample stratification for a given cell as a sample distance matrix by computing the distance between counterfactuals for all pairs of samples. More precisely, for any cell *n*, we let *D(n)* denote its sample distance matrix between counterfactuals. In this matrix, the element at the position indexed by (*s*_*a*_, *s*_*b*_), where *s*_*a*_ and *s*_*b*_ are indices representing different samples, corresponds to the Euclidean distance between the counterfactual cell states 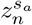 and 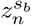.

These matrices inform sample stratification at single-cell resolution. They can first be used to identify cell populations with homogeneous sample stratifications. Clustering cells using their distance matrices as feature vectors can identify populations of cells with homogeneous sample stratifications. To do so, we embed each flattened distance matrix using PCA before clustering cells using the Leiden algorithm [62]. In any resulting cluster of cells, we then assess sample stratification in aggregate. We first compute the average sample distance matrix of the cluster, which we then use to cluster samples using hierarchical clustering.

Due to the uncertainty in *u*_*n*_, even two samples with identical underlying distributions will have non-zero distances between their estimated counterfactual cell states. To account for this uncertainty, MrVI optionally computes Monte Carlo estimates of the distribution of distances between two counterfactual cell states derived from the same sample. These distribution estimates can then be used to z-score the original distance matrix values. In detail, for a cell, *x*_*n*_, one can sample 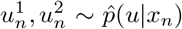, then compute the L2 distance between them, 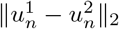. One can estimate the mean, 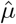, and standard deviation, 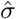, of this term with more Monte Carlo samples and use these to compute normalized distances for *D(n)* as 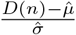.

#### A.4 Assessing compositional and expressional sample differences

With observed sample covariates at hand, MrVI can also highlight which cells have different abundances or expression levels across groups (Figure 1d). Such characteristics can, for instance, correspond to age, sex, or disease status when samples correspond to different donors. The target covariate may also be derived from a stratification of samples based on the procedure described in the previous section. This section outlines how MrVI can be employed for both DE and DA analyses assessing sample differences in gene expression and cell composition.

##### Cluster-free assessment of differences in expression

First, MrVI can characterize differential expression patterns across samples. Suppose we observe *C t*arget covariates in the form of a vector *c*^*s*^ ∈ ℝ^*C*^ for each sample *s*. To identify affected cells and genes by each target covariate, we fit the following linear model for each cell *n:*

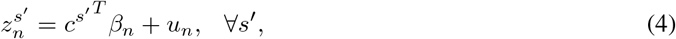

where 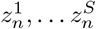 are *S c*ounterfactuals for cell *n o*btained from Equation (3). Here, *β*_*n*_ ∈ ℝ^*C×L*^ is the vector of regression coefficients obtained via least-squares regression.

##### Identifying the effect of covariates on cells

This linear model can first quantify the overall effect of an observed covariate on any cell. We compute, for any cell *n a*nd covariate index *j* ≤ *C*, the Chi-squared statistic of 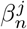. This statistic quantifies the extent to which the observed covariate *j e*xplains the variation in the counterfactual cell states.

##### Detecting cells strongly affected by covariate

The results of the linear regression can help identify cells strongly affected by a covariate. We compute the L2 norm of the vector *β*_*i*_ for a covariate *i*. This yields effect strengths in the *z r*epresentation for the specific covariates and can be used to compare effect strength across multiple cell types.

##### Detecting differentially expressed genes

The results of the linear regression can also identify DE genes associated with a given covariate in any cell. For simplicity, assume that the covariate of interest is binary. Let 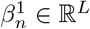 denote the regression coefficients of the covariate of interest for cell *n*. To identify the associated DE genes, we decode the counterfactual cell state 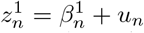 and the reference cell state 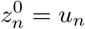. This computation yields two vectors of decoded gene expressions, denoted as 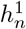 and 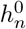. We then compute the log-fold change between these two vectors, measuring the effect of covariate *j o*n each of the observed genes *g* in the cell.

##### Accounting for out-of-distribution samples

Prior to conducting the described procedure, we first identify and discard samples that are out-of-distribution for any given cell. Samples will be out-of-distribution for a cell if no cell from that sample was collected in a similar cell state in the *u*_*n*_ space. For these samples, the model has insufficient information to accurately infer realistic counterfactual cell states in Equation (3). Thus, we conservatively discard the sample *s f*or cell state *u i*f the maximum density reached at *u w*ith respect to the approximate variational posterior distributions falls below a given threshold *τ*. More details on how the densities are computed and how *τ i*s chosen are given in Section B.

##### Assessing cluster-free differences in composition

Last, our approach can identify differentially abundant cell populations over groups of samples using log ratios of aggregated posterior densities. For this purpose, we introduce *q*_*s*_, the aggregated posterior distribution for a given sample *s*, which corresponds to 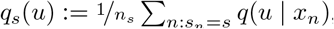, where *n*_*s*_ denotes the number of cells in the considered sample and *q(u* | *x*_*n*_) is the variational approximation to the posterior distribution over the *u s*pace for cell *n*. We can then quantify the density of any set of samples *A* ⊂ {1, … *S*} in the *u* space as 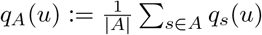.

For two disjoint sets of samples *A* and *B*, we quantify the relative overabundance of cells from *A* compared to *B* at any cell state *u* by computing the log density ratio of the aggregated posterior densities of the two groups, i.e.,

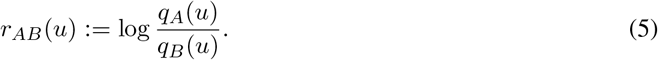

We can then identify enriched or depleted regions of *u i*n *A c*ompared to *B v*ia inspection of the log ratio *r*_*AB*_(*u)*. This approach has several benefits. As MrVI assesses differential abundance in the *u l*atent space, the captured differential abundance effects are orthogonal to the differential expression effects quantified in the previous section. Furthermore, this approach allows us to identify enriched cell states without requiring cell-type or neighborhood assignments.

At a cluster-level, we also devise a strategy to identify cell enriched or depleted subpopulations with statistical confidence. For this purpose, letting *A d*enote a cluster of interest, we collect log-ratios for (i) all cells in cluster A (ii). those not in A. We then test for difference in the mean of these two sets of log-ratios using a two-sample t-test. To avoid detecting differences as significant due to large sample sizes, the t-test rejects for the composite null that the mean difference is, in absolute value, less than a given threshold *δ*. Throughout the experiments, we set *δ =* 0.1.

### B Additional model details

#### Mixture of Gaussians prior for *u*_*n*_

MrVI posits a mixture of Gaussians (MoG) prior on *u*_*n*_, that writes as

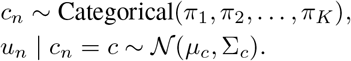

In practice, we assume the covariance matrices to be diagonal and learn *µ*_1_, …, *µ*_*K*_, Σ_1_, …, *Σ*_*K*_, and *π*_1_, …, *π*_*K*_ during training using maximum likelihood estimation.

When cell-type annotations are available, MrVI can weakly encourage the mixture of Gaussians to align with these annotations. In this case, we set *K* to be the number of unique cell-type annotations and reparameterize the mode of the Gaussian distribution as 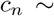 Categorical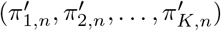, where 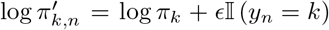. Here, *ϵ* is a positive constant (with a default of 10), *y*_*n*_ is the cell-type annotation of cell *n*, and 𝕀(.) is the indicator function.

#### Parameterization of multi-head attention layers

Two components of MrVI rely on multi-head attention layers, corresponding to the mappings *f*_*ϕ*_ and *g*_*θ*_ from Equation (1). We now provide details on how these layers are parameterized, illustrated in Figure 6.

**Figure 6:**
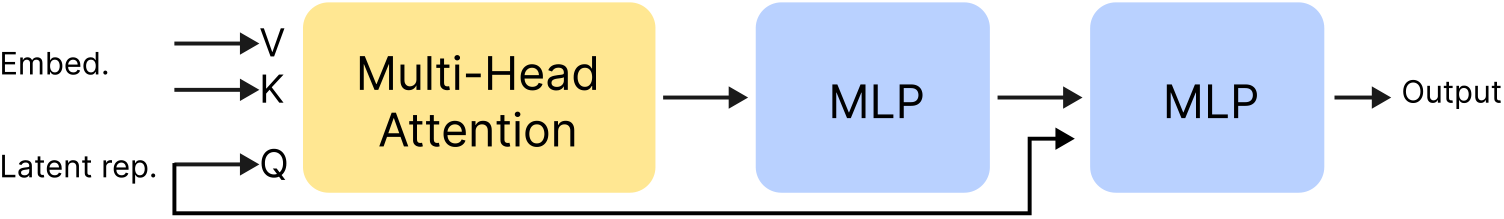
Illustration of the parameterization of the multi-head attention mechanism used in MrVI. For any cell *n*, this mechanism takes a embedding and a latent representation as inputs. These two inputs respectively serve as keys/values and queries for the attention mechanism. This output is then passed through a series of fully connected layers to obtain the final output.

**Parameterization of** *f*_*ϕ*_ *f*_*ϕ*_ takes as inputs the cell-state *u*_*n*_ and the sample *s*_*n*_ from which the cell was collected. We associate each sample ID *s* ∈ {1, … *S*} with an embedding *e*_*s*_ ∈ ℝ^*L*^, learned during training. We then rely on a multi-head attention mechanism to capture the effect of *s*_*n*_ on *u*_*n*_ in a nonlinear fashion, considering *u*_*n*_ as queries and 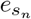 as keys/values. This output is then passed through a series of two fully connected layers, with ReLU activations, to obtain the actual output of *f*_*ϕ*_.

**Parameterization of** *g*_*θ*_ *g*_*θ*_ relies on the exact same parameterization as *f*_*ϕ*_, but takes *z*_*n*_ and *b*_*n*_ as inputs instead of *u*_*n*_ and *s*_*n*_.

#### Out-of-distribution checks

MrVI DE module actively filters for out-of-distribution cell/sample pairs. It may for instance be the case that a given sample contains no cells of a given type; in this case, this sample should be discarded for the DE analysis in Equation (4). We identify these out-of-distribution samples by considering sample-specific aggregated posterior distributions 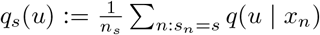. For a given cell *n*, we filter out sample *s* whenever *q*_*s*_(*u)* ≤ *τ*_*s*_, where *τ*_*s*_ is set to the 5% quantile of densities *q*_*s*_(*u)* over all cells collected in *s*. After computing the set of admissible samples for every cell, additional filtering can be performed on the level of cells to eschew those with very few admissible samples (e.g., a rare cell type observed in one sample) for which counterfactual estimates may be generally unreliable.

### C Benchmark

#### Baselines

##### Exploratory analyses

We considered two approaches that stratify samples based on differences in cell cluster abundance. Both approaches compare subcluster proportions between samples to yield distance matrices. More particularly, they subcluster each predefined cell group with the Leiden algorithm [62] using low-dimensional cell representations, i.e. PCA for Composition (PCA) or scVI for Composition (SCVI). The distance between two arbitrary samples is then defined as the Euclidean distance between their subcluster proportions.

##### Guided analyses

We also considered Milo [19] and miloDE [63], which leverage estimates for DA and DE, respectively, in guided analyses. Milo is a statistical framework that aims to detect cell neighborhoods enriched in certain sample groups based on a nearest-neighbor graph of cells. Built on top of Milo, miloDE [63] performs differential expression tests for each neighborhood identified by Milo by comparing each neighborhood against adjacent ones. These approaches, however, do not provide effect sizes for DA and DE at the cell level and instead group cells into neighborhoods that may obscure effect sizes at a single-cell resolution [18]. To compare these approaches to MrVI, we computed cell-level effect sizes for Milo and miloDE by defining cell-level effect sizes as the average effect size of the neighborhoods to which each cell belonged.

#### Metrics

##### Cell-type silhouette scores

We consider averaged silhouette width scores computed as in [31] to assess the relevance and the proper mixing of the latent representation *u u*nder the assumption that the same cell types appear across the considered samples. To do so, we first compute the silhouette score with respect to author-provided cell-type annotations. For any cell *n w*ith cell representation *r(n)*, belonging to annotation *C*_*o*_, let *d(n, C)* denote the mean distance of *r(n)* to representations of annotation *C*, excluding *n i*f *C = C*_*o*_. let *a(n)* denote the average distance of *r(n)* to cells of the same annotation, and *b(n)* the smallest mean distance of *r(n)*. The silhouette score for cell *n i*s computed as

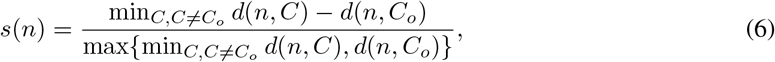

and the overall dataset silhouette score is the average of rescaled silhouette scores across all cells in the data. The rescaling, 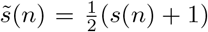, puts the dataset score in the range (0, 1) This score assesses to what extent the data representations cluster according to the annotations. When the dataset score is equal to 1, representations with the same annotation perfectly cluster together.

##### Batch silhouette scores

We also used the silhouette to measure the extent to which batch IDs mix together in the latent space. To do so, we follow the procedure described in [31], which consists of, for each previously-annotated cell type: (i) computing cell silhouette scores with respect to the batch assignments, (ii) rescaling these scores, such that 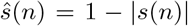, and (iii) computing an overall silhouette score computed as a weighted average of *ŝ*(*n)*, to ensure that each cell type gets the same contribution.

### D Data and preprocessing

#### Semi-synthetic experiment

We constructed a semi-synthetic dataset containing controlled DE and DA effects. Starting from a PBMC dataset of 68K cells [64], we generated a semi-synthetic scRNA-seq dataset containing a total of 32 study subjects. In particular, the original metadata has no importance in this experiment; we only relied on the gene counts to redefine synthetic notions of “cell-types”, which we will refer to as cell subsets and samples, which will here correspond to synthetic study subjects. The general recipe we employed to construct the semi-synthetic data from the original gene counts was the following. We first assigned each cell to one of five clusters, viewed as cell subsets, and then assigned study subjects to each cell in a way that introduced (1) DE between subject groups defined by *covariate 1* in cell subset A and (2) DA between subject groups defined by *covariate 2* in cell subsets B and C.

While we have no exact ground truth for the subject-subject distances, our ground truth consisted of a dendrogram, or tree, over study subjects, characterizing the similarities between subjects in terms of gene expression in the cell subset A. Specifically, all subjects sharing the same *covariate 1* value shared similar gene expression values for cells in subset A.

In two other cell subsets, denoted as B and C, cells had no differences in expression over subjects but exhibited differences in abundance, either corresponding to enrichment or depletion of these cell subsets in a specific group of samples. In particular, all subjects sharing the same *covariate 2* value shared similar proportions of cell subsets B and C but different proportions with the other subjects.

For all other cell subsets, we ensured that they neither had a DA nor a DE effect. The following are details of generating this semi-synthetic dataset.

#### Assigning cells to cell-types

To assign cells to cell subsets, we clustered cells using the Leiden algorithm [62] on the log-median normalized counts and denoted the three most abundant clusters as A, B, and C. In this experiment, it is not important for these clusters to capture plausible cell types.

#### Introducing DE between samples in cell subset A

Once cell subsets were defined, we assigned cells of subset A to study subjects to introduce DE between subjects. To achieve this, we stratified cells of subset A into subpopulations, each of which characterized a group of study subjects sharing the same expression profile for subset A. To do so, we picked 100 genes at random and performed hierarchical clustering on cells of subset A using these genes only, with a total number of eight clusters. The resulting dendrogram stratified cells of subset A into eight subpopulations, each of which contained cells that would be subsequently assigned to subjects with a shared category for *covariate 1*. To be precise, for each cluster and the associated group of subjects (each group consisting of four subjects) sharing a category for *covariate 1*, we assigned the cells uniformly at random between the four subjects. This strategy produced a total of 4 identifiers *×* 8 categories = 32 subjects.

We relied on DESeq2 [13] to compute reference LFCs for the comparison of cells of subset A in the first subject to the rest of the subjects.

#### Introducing DA between subjects in cell subsets B and C

We used a different subject assignment strategy in cell subsets B and C, specifically designed to introduce DA effects between subjects without introducing DE effects. To achieve this, we first assigned each cell of subsets B and C to one of the 32 subjects uniformly at random. Each subject was then assigned a depletion rate according to its value of *covariate 2* (one of four), which we denote *r*_*s*_, that determined the rate of over-sampling in subset B and under-sampling in subset C. In other words, if we denote the original rate of sampling for cell subsets B and C as *r*^*B*^ and *r*^*C*^, respectively, we then over-sampled cells from cell subset B in each subject *s w*ith probability *r*^*B*^ + *r*_*s*_, and under-sampled cells from cell subset C in each subject *s w*ith probability *r*^*C*^ − *r*_*s*_. This strategy ensured that the relative proportion of the other cell subsets in each subject remained constant while introducing DA effects between subjects in cell subsets B and C.

#### Subject assignments in the other cell subsets

In the remaining cell subsets, subjects were assigned uniformly at random, effectively ensuring that these cells exhibited neither DE nor DA effects.

#### COVID experiment

##### Dataset & preprocessing

The original dataset [7] contained a total of 650 thousand PBMC cells sequenced across three sites: Cambridge, Sanger, and Newcastle. We discarded cells coming from Cambridge and Sanger, and focused on data points sequenced in Newcastle. We retained the 10,000 most variable genes using Seurat v3. The resulting dataset contained 418,768 cells, originating from 55 patients. No additional cell or gene filtering was performed. All throughout the experiments, we relied on the original study annotations, that were slightly simplified to simplify the analysis according to the following scheme: B_cell ⟼ B cell, CD14 ⟼ CD14 Monocyte, CD16 ⟼ CD16 Monocyte, CD4 ⟼ CD4 T cell, CD8 ⟼ CD8 T cell, DCs ⟼ DC, gdT ⟼ gd T cell, NK_16hi ⟼ NK, NK_56hi ⟼ NK, pDC ⟼ pDC, Plasmablast ⟼ Other, Platelets ⟼ Platelet, Treg ⟼ Other, HSC ⟼ Other, MAIT ⟼ Other, Lymph_prolif ⟼ Other, RBC ⟼ Other, Mono_prolif ⟼ Other, Lymph_prolif ⟼ Other.

##### Model parameters

The sample identifier used by MrVI corresponded to `patient_id`. The batch identifier was left empty. We used the same model hyperparameters as for the sci-Plex dataset, except for the following two differences. First, we used the described mixture of Gaussians prior (*K =* 20). Second, we set the dimensions of *u a*nd *z t*o respectively be 5 and 30. We trained the model with minibatch sizes of 1,024 observations. MrVI was trained using early stopping with a patience of 30 epochs based on the validation ELBO.

##### Analysis

Visualizations of MrVI latent variables relied on minimum distortion embeddings (MDE; [65]), applied with 15 neighbors and a repulsive fraction of 0.7. We identified different profiles of sample stratifications across cells by clustering cell-specific distance matrices with Leiden (resolution= 0.*0*5), which returned three different cell clusters, one containing monocytes and dendritic cells, another one containing T cells, and the other one containing the rest of the cells. Once these clusters identified, we computed cluster-specific distance matrices by averaging cell-specific distance matrices across all cells of the cluster. We then used hierarchical clustering on these matrices to stratify donors using Ward’s method [66]. We then focused on the monocytes/dendritic cell cluster and studied DA and DE across different donor strata. We relied on Equation (5) to perform DA with MrVI. For differential expression, we compared high and low DSS patient groups, as identified in the distance matrix clustering analysis, using Equation (4). This analysis produced a cell-by-gene matrix of log fold-changes for the comparison of early to late patients, each row corresponding to a myeloid cell, and each column to a gene. We used this matrix to identify three modules of genes with potentially different patterns of DE across myeloids. For this purpose, we clustered genes by applying KMeans to the gene-by-cell matrix of log fold-changes after PCA dimensionality reduction.

#### sci-Plex experiment

##### Dataset

The sci-RNA-seq3 dataset consists of RNA expression observations from the sci-Plex chemical perturbation screen over three different cell lines (A549, MCF-7, K562) [6]. This study involved 188 small-molecule drugs, each at four different doses (10nM, 100nM, 1000nM, 10000nM), as well as untreated control samples, referred to as “vehicle” samples. The assay was conducted in 96-well plates, with two biological replicates for each drug-dose combination. We chose to concatenate the drug name and dose level as the modeled target covariate and the plate number as the modeled nuisance covariate. So, the two biological replicates mapped to each drug-dose combination were treated as one sample. Because each pair of biological replicates was conducted on different plates, any significant differences between the replicates were corrected for in the *z*_*n*_ latent space. The naming convention used in the figures for the sample covariate is `“{drug name}_{dosage (nM)}”`.

##### Preprocessing

Several steps were taken to preprocess the dataset. As discussed in [6], many of the small-molecule perturbations tested had little to no effect on the cell lines. To focus the analysis on the perturbations with significant effect, we performed a simple differential gene expression (DEG) analysis to filter out a subset of drugs with no effect and to simplify visualizations later for each cell line. For each cell line, we performed a t-test-based differential expression test for each drug-dose combination with respect to the vehicle in each cell line and adjusted the p-values with Benjamini-Hochberg correction. We defined a DEG as one with a p-value less than 0.*0*5 and an absolute log-fold change (LFC) greater than 0.*5*. Then, we created a histogram of the number of DEGs for the maximum dose (10000nM) of each drug-cell-line combination (**Figure S7**). We chose 3, *0*00 DEGs as a cutoff to capture the tail of this histogram, which should generally correspond to the combinations with significant, widespread differences from the vehicle cells. For the remaining analysis, we filtered out drugs that did not reach this cutoff for any dose-cell-line combination. Additionally, we tracked which drug-dose combinations reached the cutoff number of DEGs for each cell line for visualization purposes. The resulting dataset combining all three cell lines contained 251,088 cells with 92 drugs at all four doses and the vehicle cells. Last, we applied highly variable gene (HVG) selection using Seurat v3 [67] with the cell line as the batch key and retained the top 5, *0*00 HVGs.

##### Model parameters

MrVI was run separately on each of the three cell lines with the same set of hyperparameters. Since only one cell line was contained in the dataset for each model fit, we expected *u*_*n*_ to contain continuous variation corresponding to cell cycling effects but otherwise lack any variation corresponding to inter-cell-line differences. For this reason, a mixture of Gaussians prior was not used for *u*_*n*_ in these experiments. Instead, we assumed *u*_*n*_ to follow an isotropic Gaussian distribution (*K =* 1 with 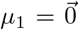 and Σ_1_ as the identity matrix in Equation (1)). Otherwise, the model parameters mostly follow our recommended defaults, including the use of the MAP estimate on *z*_*n*_|*u*_*n*_, an isotropic Normal prior on *z*_*n*_ − *u*_*n*_, and attention-based decoders. The dimensions of the *u*_*n*_ and *z*_*n*_ latent spaces were the main hyperparameters that needed to be tuned accordingly. To ensure the model fit reflected prior knowledge about the sci-Plex dataset, we developed custom metrics to select the final hyperparameters. We discuss this in the following section.

##### Model selection

We developed two dataset-specific metrics to determine our final model hyperparameters. First, we created a metric to reflect how similar the drug-dose combinations using the same drug perturbation were according to the model. Generally, we expected the effects of different dosages of the same drug to be more similar than the effects of two entirely different drugs. To capture this pattern for a given model fit, we computed the average percentile of the sample distances between same-drug-different-dose sample pairs relative to all pairwise sample distances (excluding distances to self). A lower value for this metric validated that the model was able to capture this general pattern without access to the drug-dose metadata. Second, we used L1000 bulk gene expression data [37] via the iLINCS platform [68] to validate drug-drug similarities determined by the models. Data for a subset of small molecules were available for the A549 and MCF7 cell lines. For each small molecule, we determined the set of DEGs using the criteria of p-values ≤ 0.*0*5 and absolute log-fold change ≥ 0.*5* using the differential expression data provided in the dataset. Then, we constructed a drug-drug similarity matrix for each cell line by computing the Jaccard similarity between the DEG sets of each drug. The Jaccard similarity is defined as 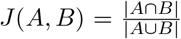 for gene sets *A, B*. We clustered the drugs using the Leiden algorithm over the similarity matrix. Finally, we computed the rescaled silhouette score, 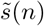 (Equation 6), of these clusters with respect to the subset of the sample distance matrix output by the MrVI model. In this case, a higher score implied the model’s sample distance matrix aligned more closely with the results of the L1000 gene expression profiles.

Lastly, we looked at the validation ELBO for each model, which describes how well the model generalizes to data it has not trained on. We found that optimizing over the validation ELBO alone did not provide good sample stratification according to the other metrics. This may be because, for smaller latent dimensions, the parameter space is smaller and thus can achieve a better ELBO by Bayesian Occam’s Razor [69]. Effectively, these smaller models tradeoff, modeling finer patterns in sample-specific variation for lower penalty attributed to the model prior. The other metrics provide a way to decide between the models with the best validation ELBO scores.

We tested different values for the dimensions of dim(*u*_*n*_) ∈ [2, *5, 1*0, *3*0, *5*0] and dim(*z*_*n*_) ∈ [2, *5, 1*0, *3*0, *5*0] where dim(*u*_*n*_) ≤ dim(*z*_*n*_) and found dim(*u*_*n*_) = 10 and dim(*z*_*n*_) = 30 had top values for all of the described metrics (**Figure S8**).

##### Analysis

First, as a high-level check to see that *u*_*n*_ and *z*_*n*_ exhibited the expected variation, we visualized each space using a minimum distortion embedding (MDE) with a repulsive fraction of 1.5 and 15 neighbors. We colored the projection by cell cycle and drug class (as reported in the original dataset) (**Figure 4a,b**). The cell-cycle labels were computed using ScanPy’s `tl.score_genes_cell_cycle` function and the cell-cycle markers from [70].

Then, we computed the sample distance matrices and took the mean across all cells. Typically, one would perform the aggregation for subsets of cells exhibiting similar sample stratification (e.g., cell types). However, in this case, since each model was run over a single, relatively homogeneous cell type, the aggregation was performed over all cells. To ensure this aggregation was reasonable in a data-driven manner, we ran PCA over the learned sample distance matrices to check if there were groups of cells with distinct stratification (**Figure 4c**). We found that the top two PCs (*> 4*0% of variation explained for each cell line) showed one cluster of cells for every cell line.

Then, we visualized the distance matrices reported by each model. To focus each visualization on the drugs with the most significant effects on each cell line, we filtered for the drug-dose-cell-line combinations in the top 20 percent based on distance from the vehicle controls. For example, this left 75 drug-dose combinations, including the vehicle, for the A549 cell line. Over this filtered matrix, we performed hierarchical clustering using Ward’s method. We then chose the number of clusters as roughly the elbow of the plot between the sum of squared differences and the number of clusters (**Figure S12**).

For each cluster found by the hierarchical clustering, we applied our model-based differential expression procedure for each cluster against the vehicle controls for the appropriate cell line. For each gene and each cluster, we averaged all the log-fold changes (**Figure S13**) then defined the DEGs as those exceeding an absolute average log-fold change of 1. We applied gene set enrichment analysis (GSEA) [40] using the MSigDB Hallmark 2020 gene set [41] separately over the set of up-regulated genes (genes with average LFC *> 1*) and the set of down-regulated genes (genes with average LFC *< 1*) and visualized the resulting scores for MSigDB gene sets where at least one cluster had a p-value *< 0*.*0*5 (**Figure 4f**).

We presented the results of the A549 cell line in the main paper (**Figure 4**) and left the results of the remaining two cell lines, MCF7 and K562, to the supplement (**Figures S14**, **S15**). We include the analogous supplementary figures for MCF7 (**Figures S16**, **S17**, **S18**, **S19**) and K562 (**Figures S20**, **S21**, **S22**, **S23**) as well.

#### IBD experiment

##### Dataset & preprocessing

We downloaded the data set from the Broad Single Cell Portal and concatenated the data sets in all organs and fractions. As the original dataset did not contain raw counts, we reverted the applied normalization in the following way. We transformed the normalized counts by expm1, divided them by 100,000 (original normalization), and multiplied by the original counts per cell. As the resulting gene expressions still contained small deviations from expected integer representations due to numerical inaccuracies, we rounded the values to the closest integer. We selected 10,000 highly variable genes using `“seurat_v3”` flavor in scanpy and using the respective 10X chemistry as batch key. We also filtered cells expressing more than 300 genes, 1K counts, or more than 20 mitochondrial reads, as these cells contained low-quality events.

##### Model parameters

We used a cell-type prior for the mixture of Gaussian, set the dimensionality of z-space to 200 and of u-space to 10 and reduced n_epochs_kl_warmup to 25. For all other hyperparameters, we used default parameters. We set the biosample_id as the sample key and the concatenation of layer and chemistry as the batch key and trained the resulting model for 150 epochs.

##### Analysis

For the UMAP embedding of all cells, we used 7 nearest neighbors and a minimum distance of 0.3. Sample distances were computed with normalized distances and using 20 Monte-Carlo samples. We used the upper-triangle of these distances and flattened those, followed by scaling these values and computed 50 PCs on the distance matrices. The resulting PCs were used for Leiden clustering (10 neighbors, cosine distance) with a resolution of 0.1 to yield the coarse cell-type labels displayed in **Figure 5a**. We annotated these coarse labels on the basis of the composition of the original labels in each coarse label.

The sample similarity matrices were displayed using PyComplexHeatmap. We subset the distance matrix to each coarse cell type and compute the mean distance. Furthermore, we filter all samples with an average admissibility below 0.3 using the *ball* criterion with a quantile threshold of 0.05. The linkage and clusters were computed using scipy.cluster.hierarchy with fcluster setting it to 5 clusters and using the criterion *maxclust*. In PyComplexHeatmap we used ClusterMapPlotter without scaling and without row or column clustering.

To study sample stratification, we performed a multivariate analysis with batch-specific offsets, filtering donors with a quantile of 0.0, and an L1 regularization of 0.01. We computed 50 mc samples and computed differentially expressed genes for each cluster identified from the sample distances, using the same approach as in the COVID experiment.

For further analysis, we excluded samples collected during surgery, as the experimental design of this study was rather imbalanced; the batch of surgical samples only included patients with stenosis, while the endoscopy biopsies contained a mix of patients with different disease subtypes. This imbalance introduces biases in downstream analysis because we found that surgical samples had significantly fewer immune cells and more stromal cells than biopsy samples (**Figures 5a**). To study stenosis, we reran the multivariate analysis on all cells from diseased individuals (excluding healthy controls and samples from surgeries) using Type (inflamed, non-inflamed), Site (colon, ileum), and Sex and Disease Behavior (B1, B2, B3) as covariates. We computed the DE genes as described in the previous paragraph and displayed the effect sizes of the multivariate analysis to identify cell populations strongly affected by covariates. For the UMAP plots of stromal cells, we used 20 nearest neighbors and a minimum distance of 0.3. We relied on Equation (5) to perform DA with MrVI. We filtered out all samples with less than 50 stromal cells as the DA estimates become noisy for low numbers of cells. Consequently, the Epithelial layer samples, which contained few stromal cells, were filtered out, leaving only the Lamina propria samples. To obtain a smooth estimate of differential gene expression, we compute normalized KNN connectivities (20 neighbors, normalized to a sum of 1) and multiply those normalized connectivities twice with the estimated log-fold changes. For violin plots and dotplots, we used the defaults in Scanpy. For the dotplots, we selected the top three marker genes in Scanpy using default setting for DE testing based on raw expression.

## Code availability and reproducibility

MrVI is available within scvi-tools (https://scvi-tools.org/). Code to reproduce the experiments is available at https://github.com/YosefLab/mrvi-reproducibility.

## Supplementary Information

**Figure S1:**
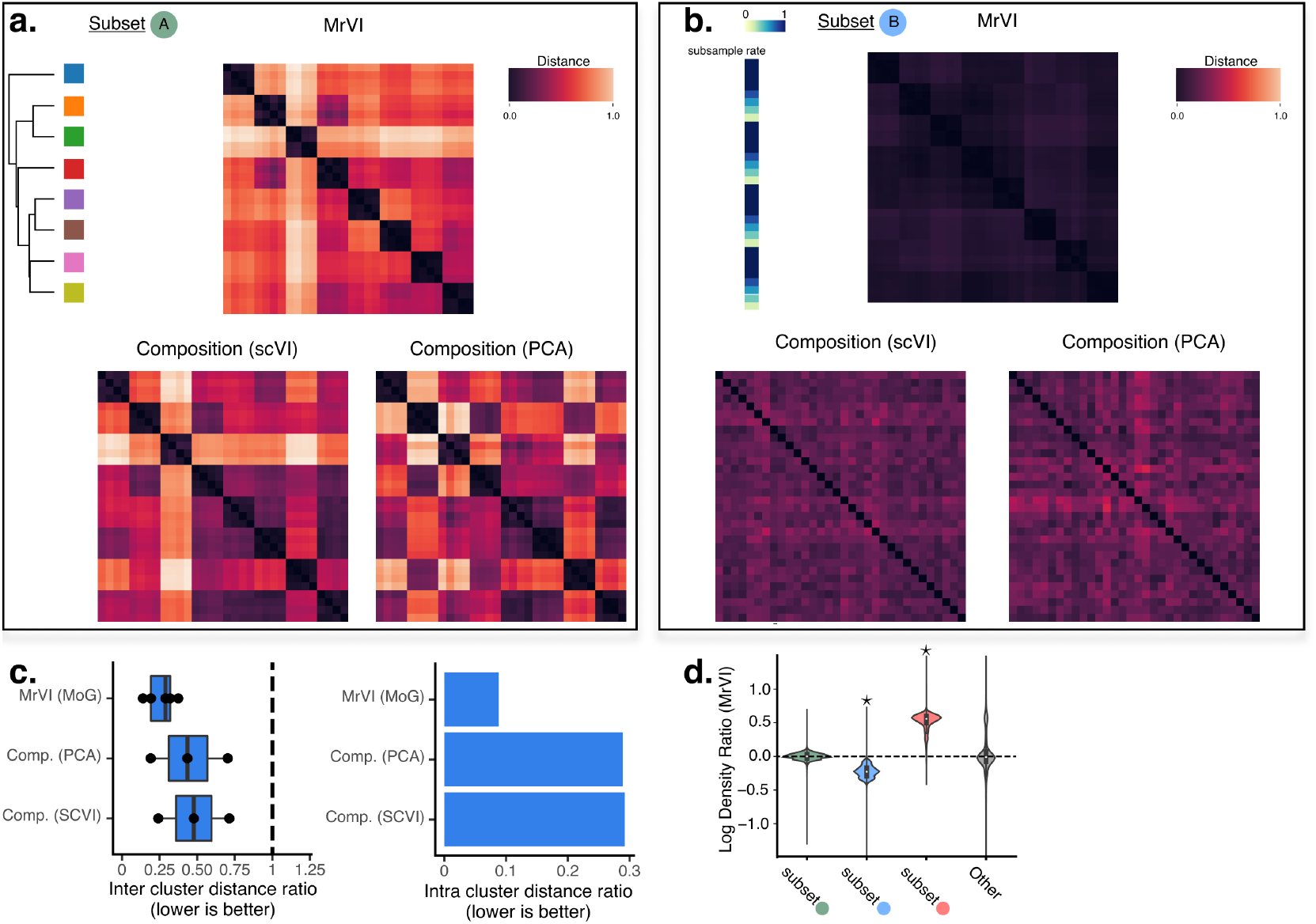
Supplementary results for the semi-synthetic experiment. **a**. and **b**. Distance matrices for MrVI and for the compositional baselines, respectively computed for cells containing DE and DA sample effects. The ordering of rows and columns is the same for all approaches. Because cell of subset Ⓑ were randomly assigned to synthetic samples, we do not expect to see any differences between samples in b. **c**. Distance matrix quality scores for the semi-synthetic experiment. *Left*: Distributions of mean distance ratio in sample-distance matrix between cell subset A and all other cell subsets compared for different methods (*lower is better*). For every cell cluster *r* different than A (the cluster containing DE effects), we computed the ratio of the average elements of the distance matrix for *r* to the average elements of the distance matrix for A. Elements of the sample distance matrix corresponding to the distance of the sample to itself were excluded for the computation of the average. Since in *r* there is no intra-cell-type variation by design, we expect these ratios to be as close to 0 as possible. The plot shows the distribution of these ratios across all *r*. MrVI produces significantly smaller distance ratios than the compositional approaches (Mann-Whitney U test, *p <* 0.1). *Right*: Mean ratio of within-category smallest distance over between-category smallest distance for *covariate 1* across different methods (*lower is better*). We calculated, for every row of the matrix associated with cells in cluster *A*, the ratio of the average distance of the blocks belonging to the same synthetic subjects to the average distance of the blocks belonging to different synthetic subjects. Displayed is the average over all rows, which we denote as intra cluster distance ratio. **d**. Violin plot of MrVI DA log ratios displayed in Figure 2d. Stars denote statistically significant DA(*p <* 10^−8^) in the given subset based on t-tests and after Benjamini-Hochberg correction [71] (see **Methods** for more details). In particular, only the two cell subsets that are DA by construction are detected DA.

**Figure S2:**
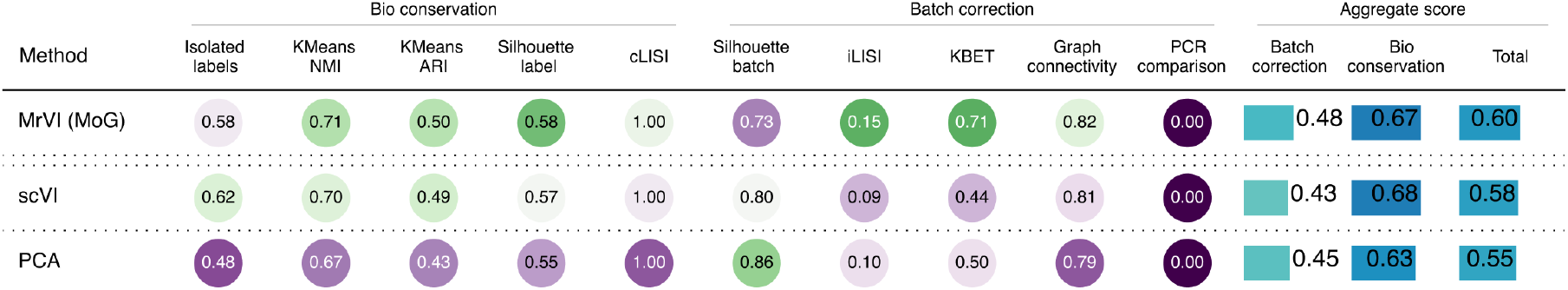
SCIB metrics for MrVI, PCA, and scVI evaluated on the COVID-19 experiment. Here, scVI used donor IDs as batch indices, and PCA was applied with *K* = 50 first principal components on log-counts per 10^4^ normalized data. We computed the SCIB metrics using cell-type annotations (**3C**) from the original study as labels, and the donor ID as the batch key.

**Figure S3:**
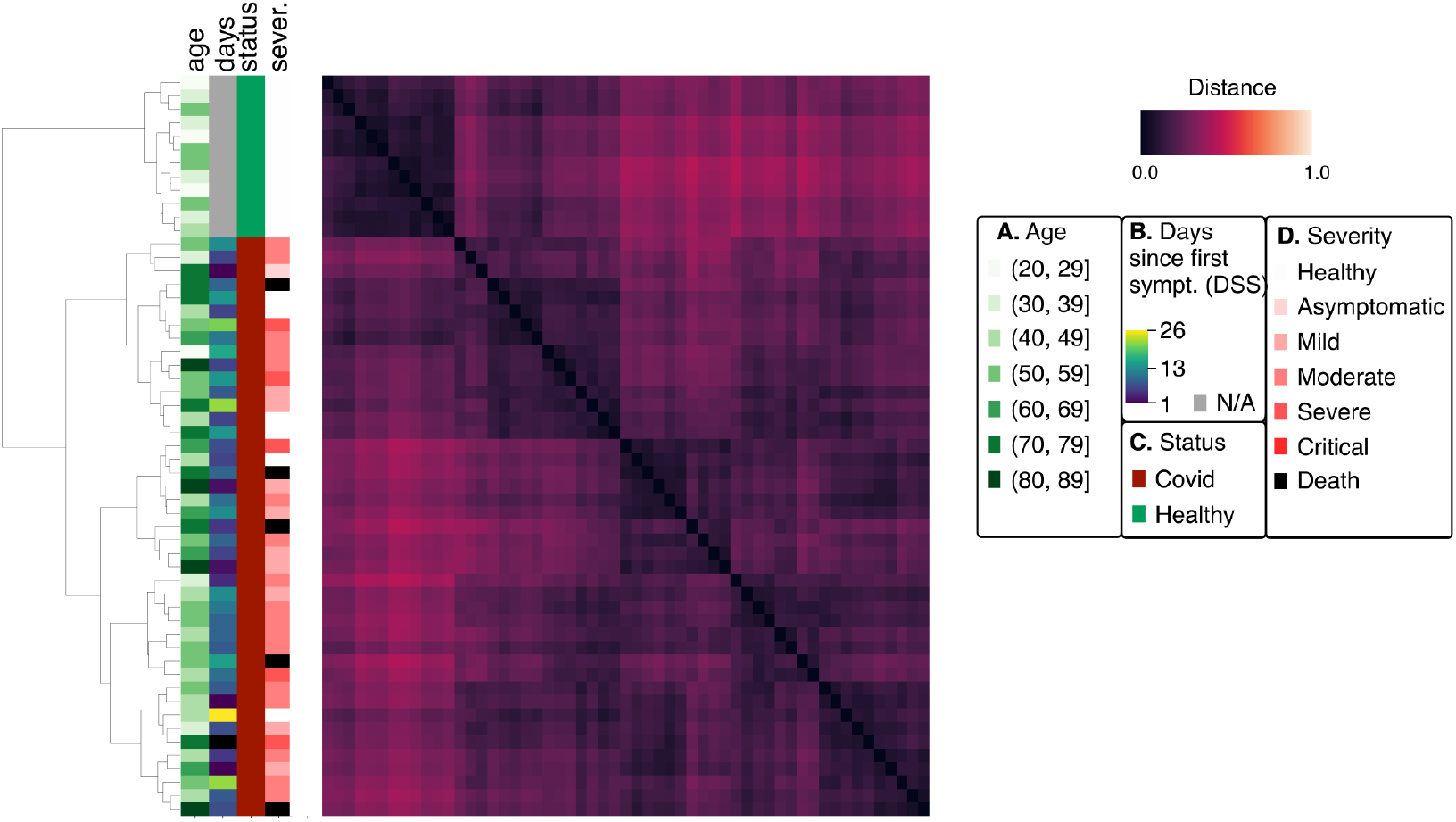
Distance matrix averaged over all cells for cluster Ⓒ, corresponding to B cells, identified in Figure 3**c** for the COVID-19 experiment. This figure relies on the same color scheme as in Figure 3**d**. We see a clear separation of healthy individuals.

**Figure S4:**
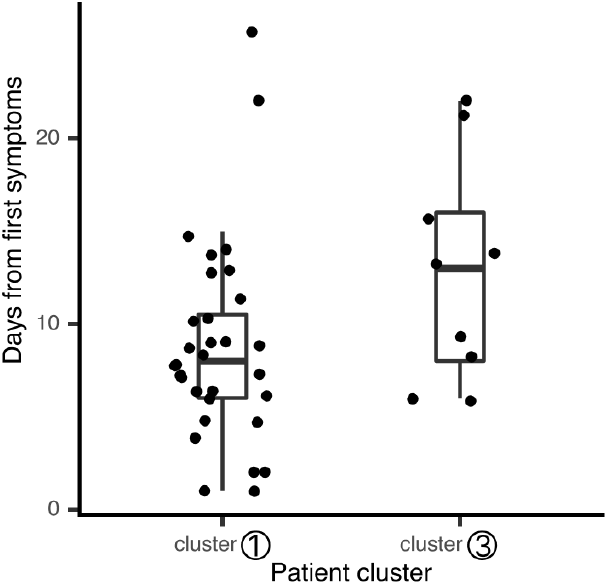
Comparison of time since symptom onset (“Days from onset” in [7]) for the two COVID-19 patient clusters from Figure 3**d**. These differences are significant under a Mann-Whitney U test (*p <* 0.05). We find several outliers, which might be attributed to the exact course of disease, or to noise due to the fact that symptom onset is self-reported. This ambiguity can’t be resolved with the current data.

**Figure S5:**
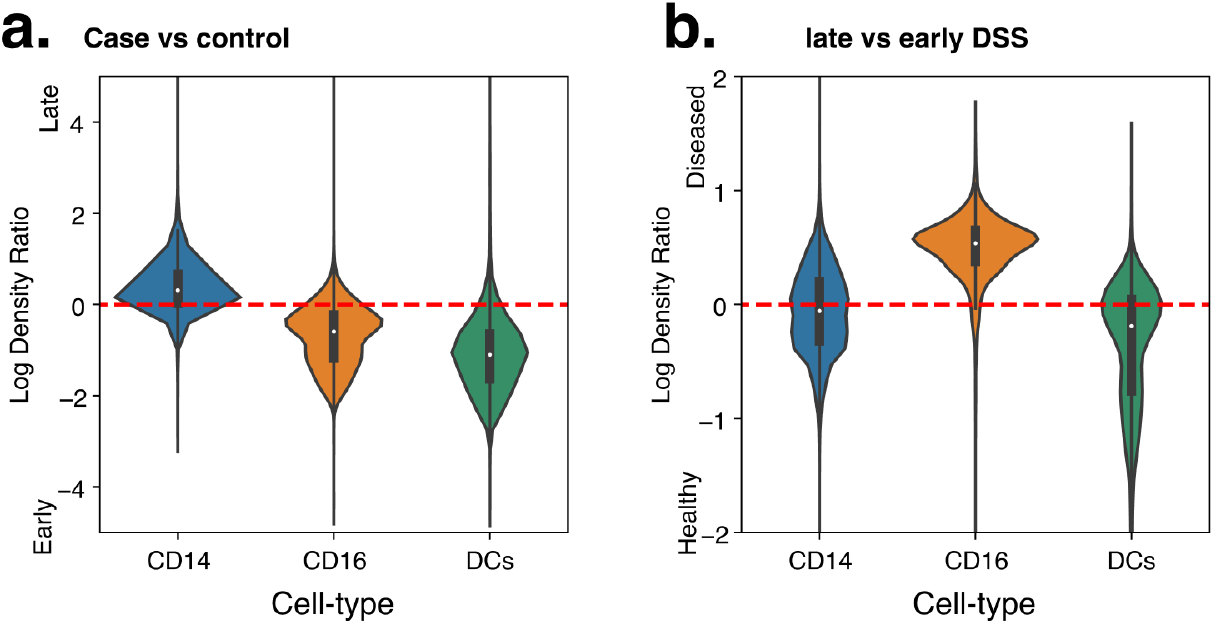
Comparison of the log density ratio displayed in Figure 3e across the different cell types contained in cluster Ⓐ.

**Figure S6:**
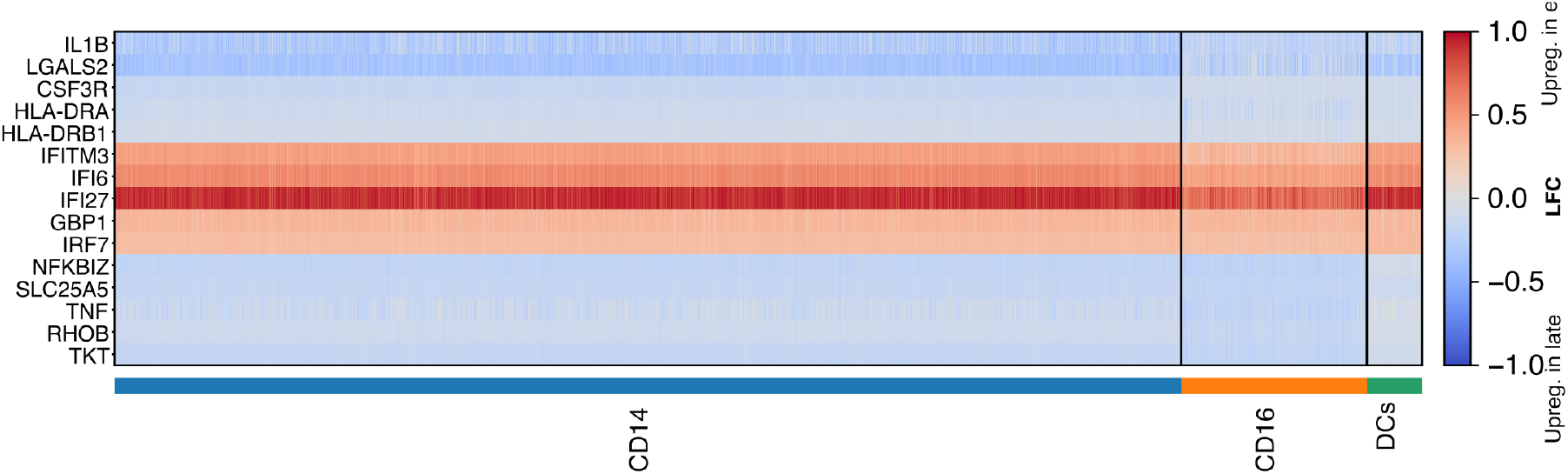
Heatmap of MrVI LFCs for the DE genes identified by in Figure 3f, for the comparison of the two groups of COVID patients, corresponding to late and early onsets.

**Figure S7:**
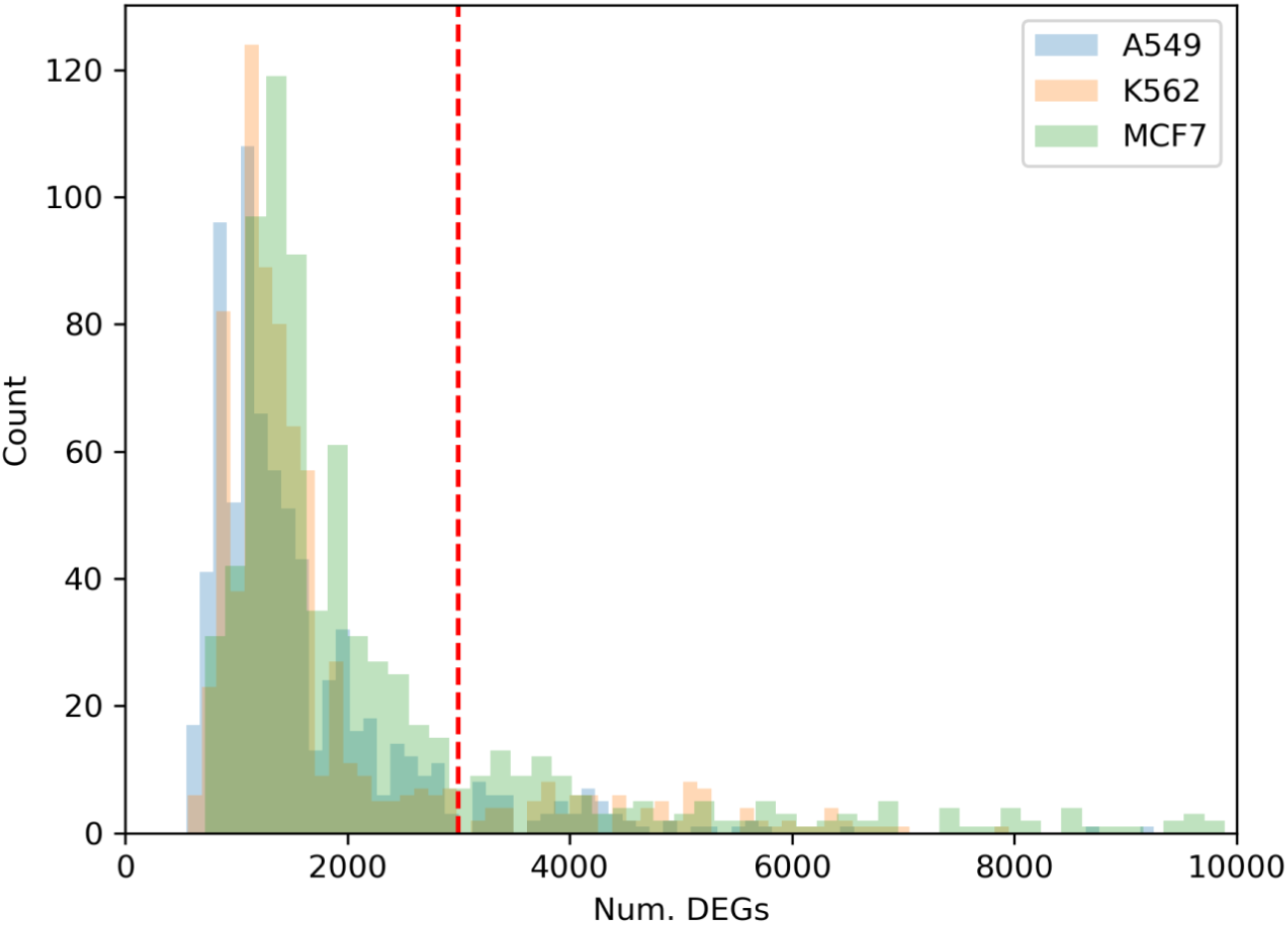
Histograms of the number of DEGs for the maximum dose (10000nM) of each drug-cell-line combination with respect to the vehicle for the three cell lines in the sci-Plex dataset. The vertical red dotted line corresponds to the cutoff used to filter out drugs with insignificant effects in all of the cell lines.

**Figure S8:**
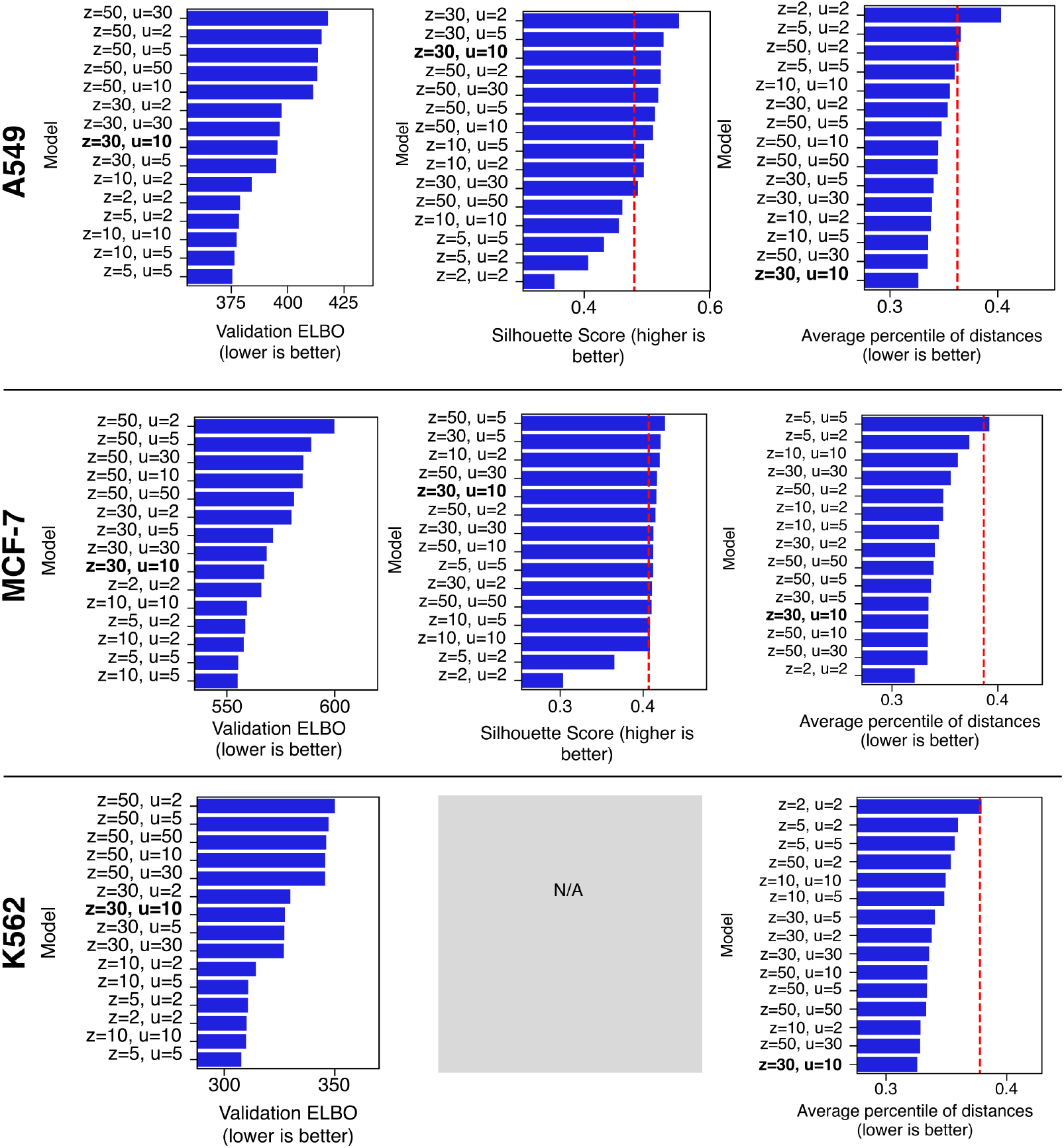
Comparison of performance of MrVI for different parameter combinations for the sci-Plex dataset. Model ELBO evaluated on validation cells, silhouette score with respect to clusters found in transcriptomic Connectivity Map data, and average percentile of within-drug sample distances over each cell line. There was no available Connectivity Map data for the K562 cell line, so we could not compute the silhouette metric for this cell line. The bold configuration denotes the hyperparameters chosen for the analysis in Figure 4, S14, S15 respectively. The red dotted vertical lines indicate the best performance between the CompositionPCA and CompositionSCVI baselines for the silhouette score and average percentile metrics.

**Figure S9:**
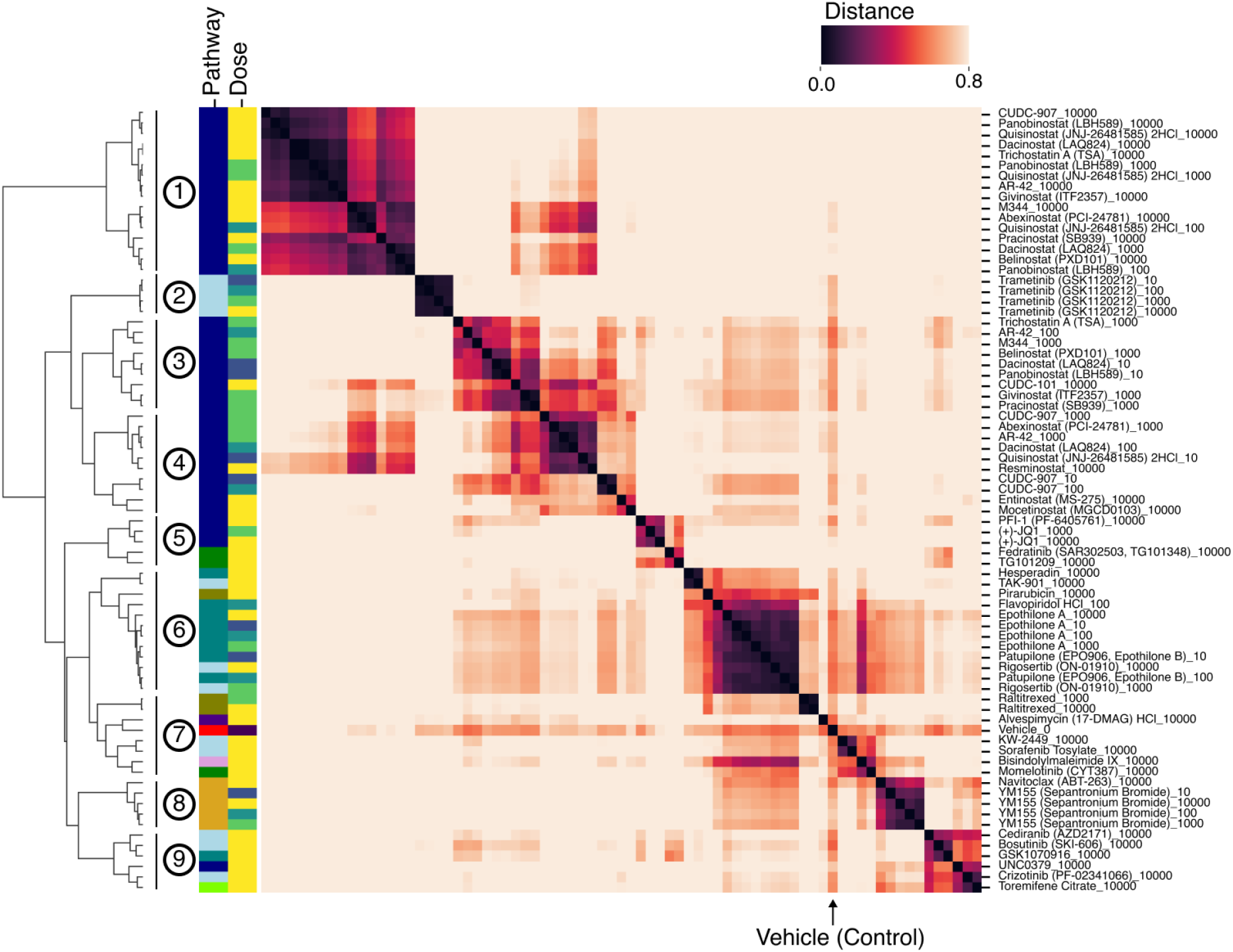
Copy of Figure 4e with row labels for each drug-dose combination.

**Figure S10:**
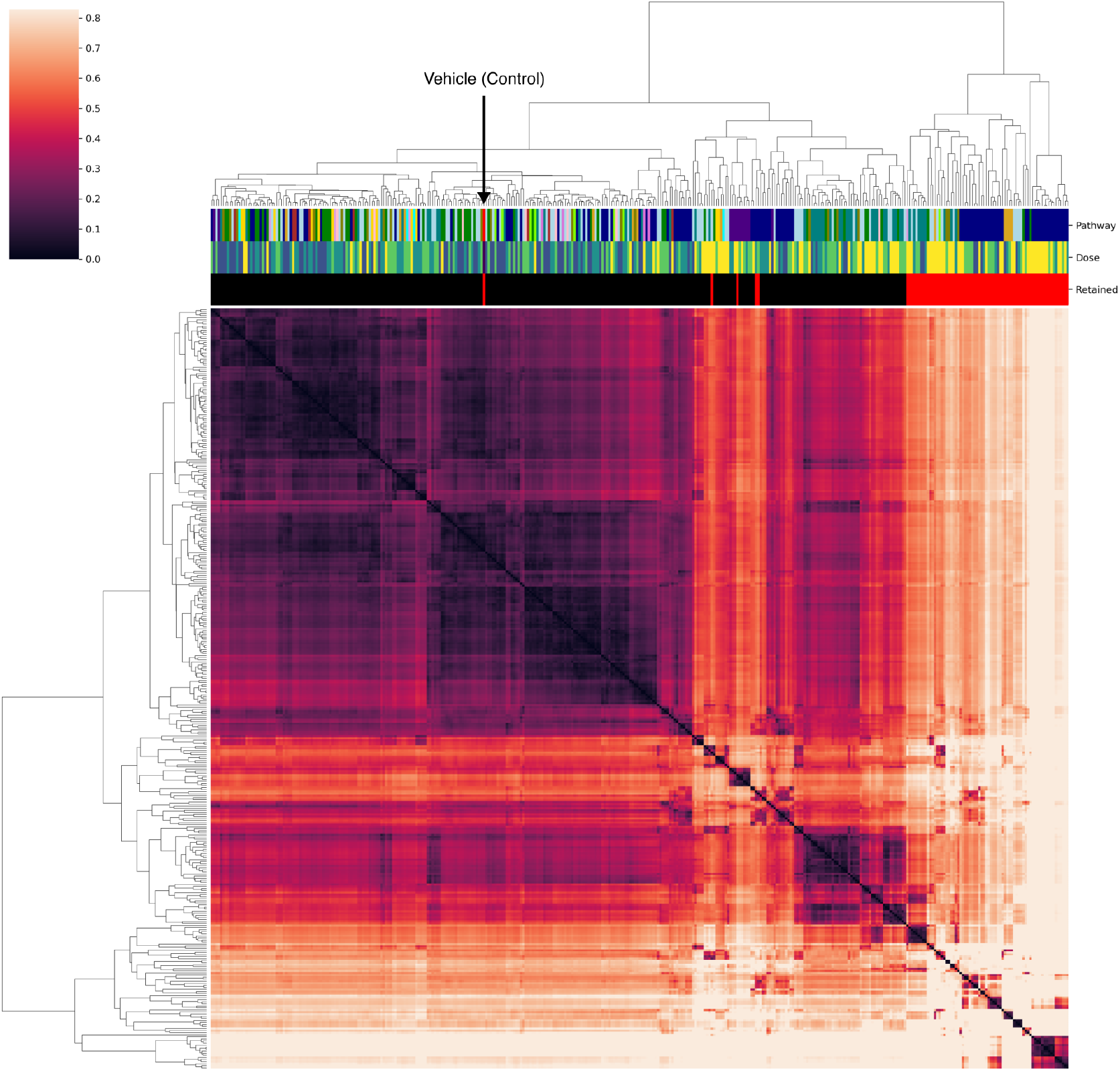
Sample distances of all 369 samples (92 drugs at four doses and vehicle) used in the analysis for the A549 cell line. The columns are annotated by each drug’s pathway annotation from the original study, dosage level, and whether the sample was retained for the remaining analysis (top 20 percent of samples based on distance from vehicle). The hierarchical clustering was performed with the Ward variance minimization algorithm.

**Figure S11:**
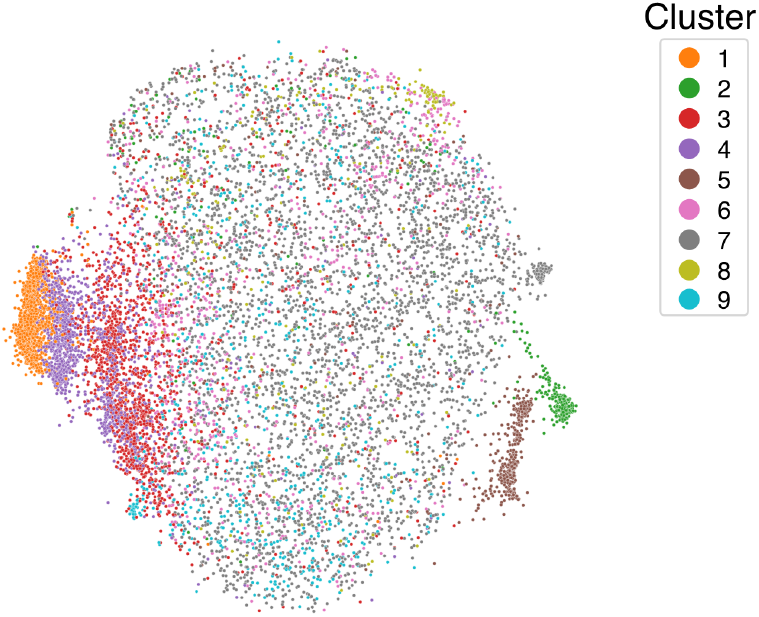
MDE of the *z* latent space from MrVI of data from the A549 cell line colored by the same cluster assignments as in Figure 4e. Clusters ➂, ➃, ➀ capture a large group of samples with rityr and increasingly divergent cell states corresponding to increasingly larger doses of HDAC inhibitors.

**Figure S12:**
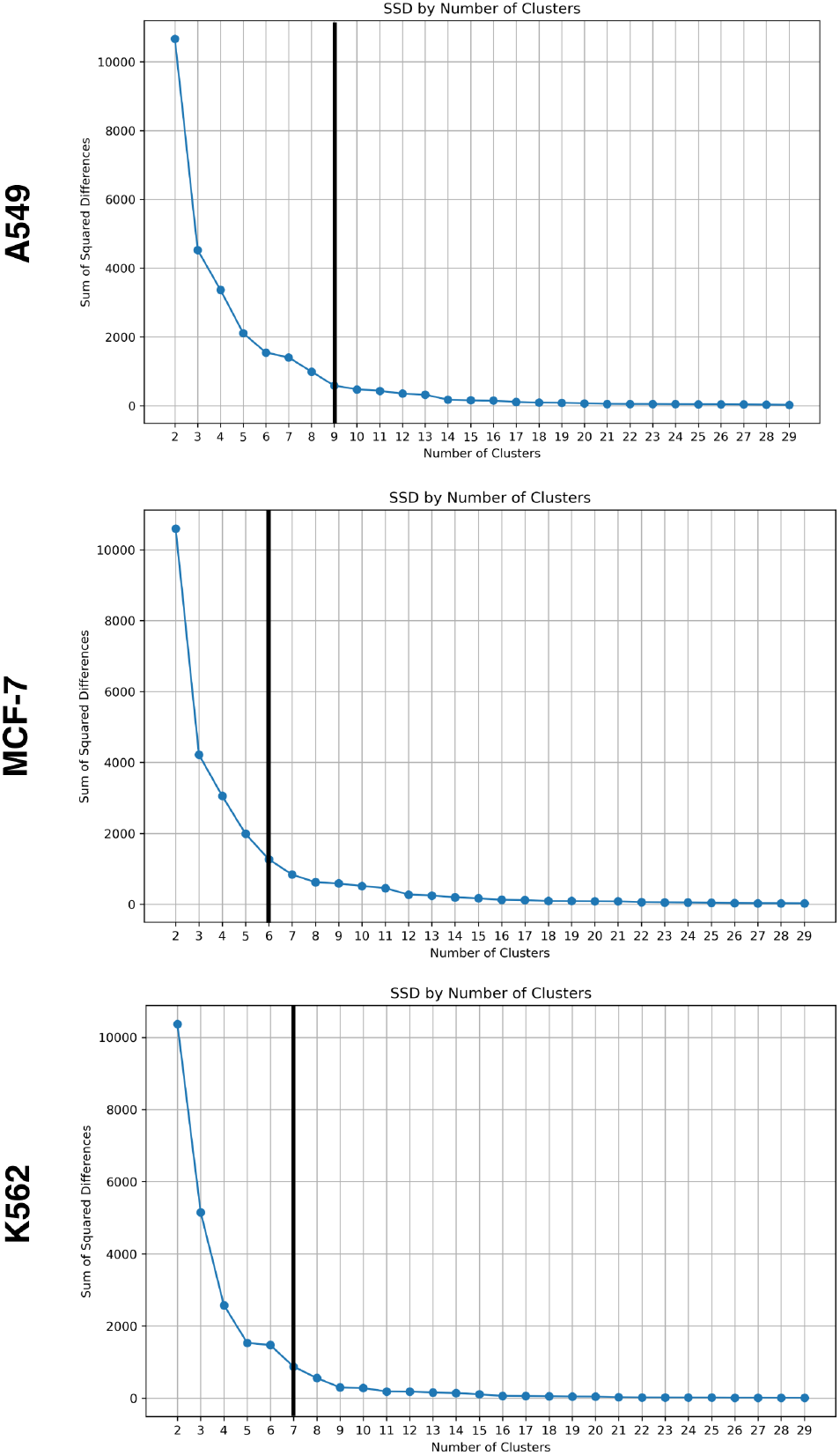
Sum of squared differences against the number of clusters used for hierarchical clustering for each cell line in the sci-Plex dataset. This metric was computed by applying the `maxclust` algorithm in `scipy.cluster.hierarchy.fcluster` over the output of agglomerative clustering, then aggregating the within-cluster, pairwise squared differences. Each vertical line denotes the number of clusters selected for the remaining analysis for each cell line, which was identified as the elbow point from a visual inspection of the plot.

**Figure S13:**
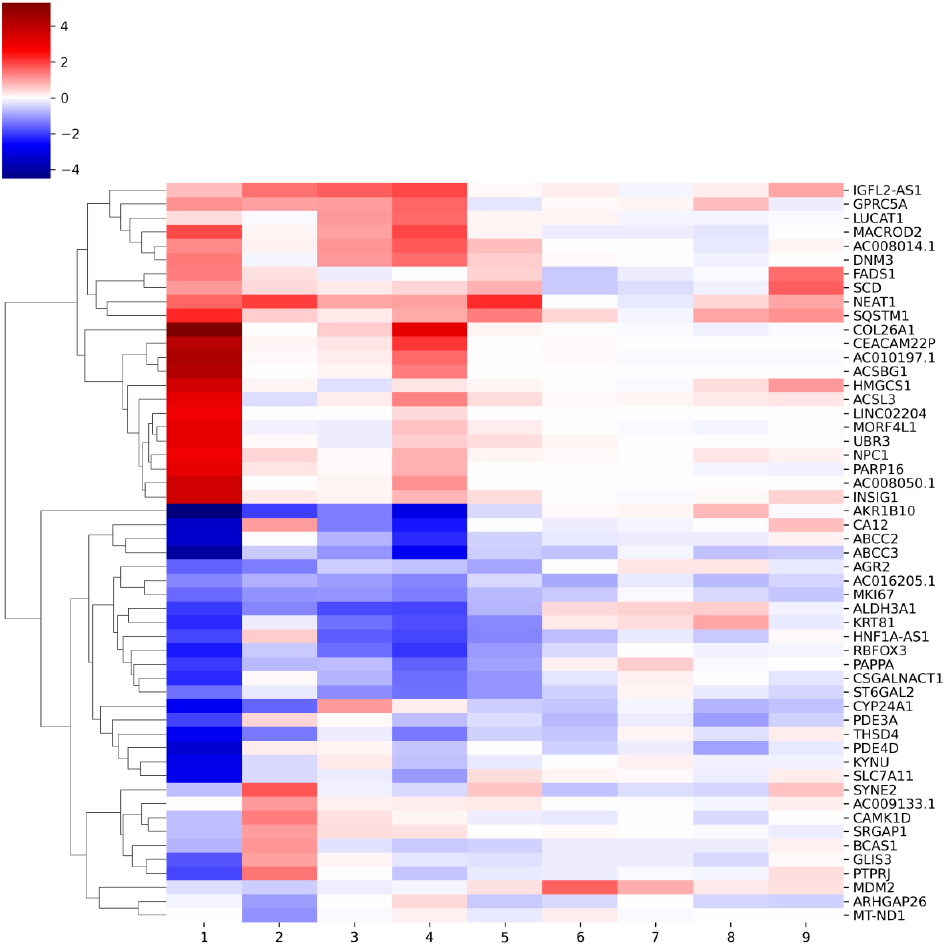
Heatmap of LFCs of the top DE genes identified by MrVI in the A549 cell line. Each column corresponds to a cluster as labeled in Figure 4e. Displayed is the LFC estimated by MrVI averaged over all cells. Concordant with the distance matrix, we find the most significant and widespread gene-specific effects in clusters ➀ and ➃.

**Figure S14:**
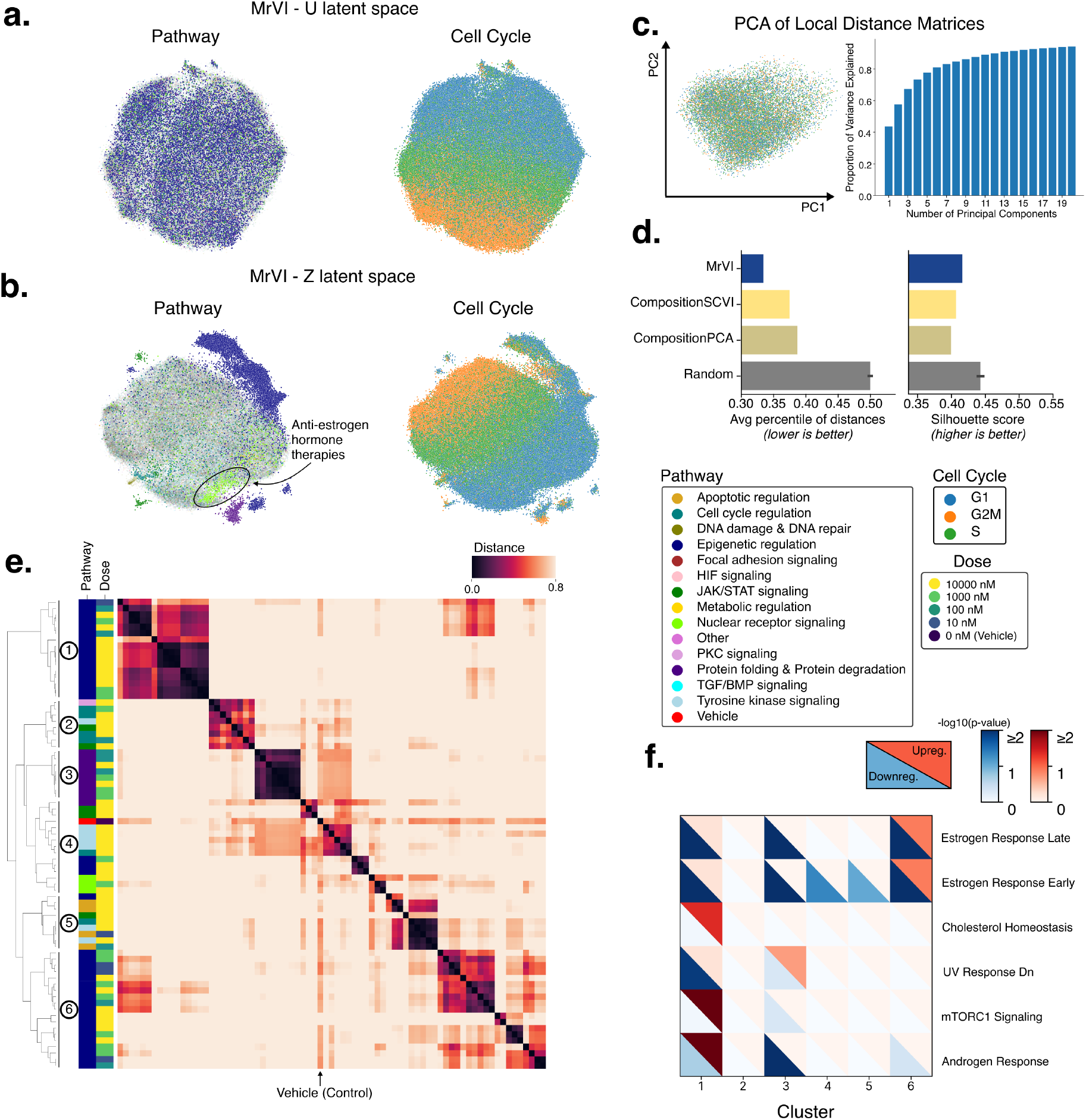
Analysis of the MCF-7 cell line in the sci-Plex experiment. MrVI was fit over 92 drugs each at four doses that passed the DE-gene filter. The analysis is performed in a way similar to **4. a. and b**. MDEs of the *u* and *z* latent spaces from MrVI colored by the target pathway of the drug used to treat each cell (*left*) and the cell cycle stage of each cell (*right*). For the MDEs colored by target pathway, only the top 20 percent of samples based on the distance from the vehicle are shown in full opacity. **c**. PCA of sample distance matrices. *left*: Scatterplot of all local sample distance matrices projected onto the top two principal components colored by cell cycle stage displays no visual subclusters. *right*: Bar plot of the proportion of variance explained against the number of principal components used. **d**. Barplots comparing MrVI against the benchmark methods for two performance metrics that determine alignment with prior knowledge. (left) The average percentile of distances measures how much closer samples with the same drug and different doses are to each other relative to the rest of the distances. We expect the average percentile to be low. (right) Silhouette score of sample clusters with similarities inferred from DEG sets in the Connectivity Map dataset. This metric measures whether the clusters are consistent. **e**. Hierarchically clustered sample distance matrix. The rows of the distance matrix are annotated by the pathway and dose of the respective sample (drug-dose combination) and by the clusters inferred from the distance matrix. For figures (e) and (f), the analysis is performed over the top 20 percent of drug-dose combinations (74 out of 368) based on their distance from the vehicle. **f**. A heatmap of Gene Set Enrichment Analysis (GSEA) scores for the Human MSigDB Hallmark gene set collection for differentially expressed genes found for each cluster found in panel (e) with respect to the vehicle cells. The upper-right triangle of each tile represents the score for the set of up-regulated DE genes, and the bottom-left triangle represents the score for the set of down-regulated DE genes.

**Figure S15:**
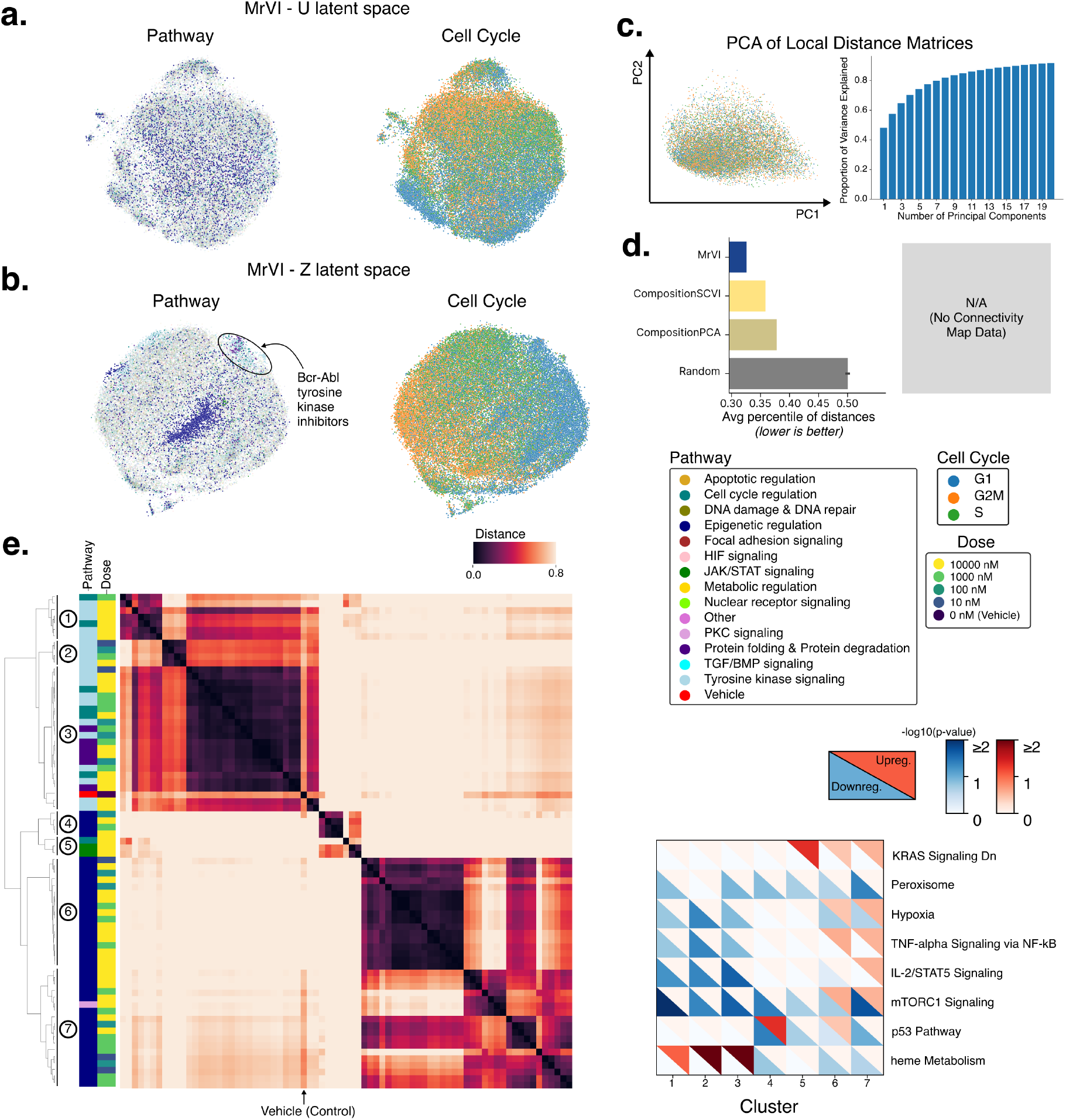
Analysis of the K562 cell line in the sci-Plex experiment. The analysis is performed in a way similar to **4**. MrVI was fit over 92 drugs each at four doses that passed our simple DE-gene filter. **a. and b**. MDEs of the *u* and *z* latent spaces from MrVI colored by the target pathway of the drug used to treat each cell (*left*) and the cell cycle stage of each cell (*right*). For the MDEs colored by target pathway, only the top 20 percent of samples based on the distance from the vehicle are shown in full opacity. **c**. PCA of sample distance matrices. *Left*: Scatterplot of all local sample distance matrices projected onto the top two principal components colored by cell cycle stage displays no visual subclusters. *Right*: Bar plot of the proportion of variance explained against the number of principal components used. **d**. Barplot comparing MrVI against the benchmark methods for a performance metric that determines alignment with prior knowledge. The average percentile of distances measures how much closer samples with the same drug and different doses are to each other relative to the rest of the distances. We expect the average percentile to be low. There was no available Connectivity Map data for the K562 cell line, so we could not compute the silhouette metric for this dataset. **e**. Hierarchically clustered sample distance matrix. The rows of the distance matrix are annotated by the pathway and dose of the respective sample (drug-dose combination) and by the clusters inferred from the distance matrix. For figures (e) and (f), the analysis is performed over the top 20 percent of drug-dose combinations (74 out of 368) based on their distance from the vehicle. **f**. A heatmap of Gene Set Enrichment Analysis (GSEA) scores for the Human MSigDB Hallmark gene set collection for differentially expressed genes found for each cluster found in panel (e) with respect to the vehicle cells. The upper-right triangle of each tile represents the score for the set of up-regulated DE genes, and the bottom-left triangle represents the score for the set of down-regulated DE genes.

**Figure S16:**
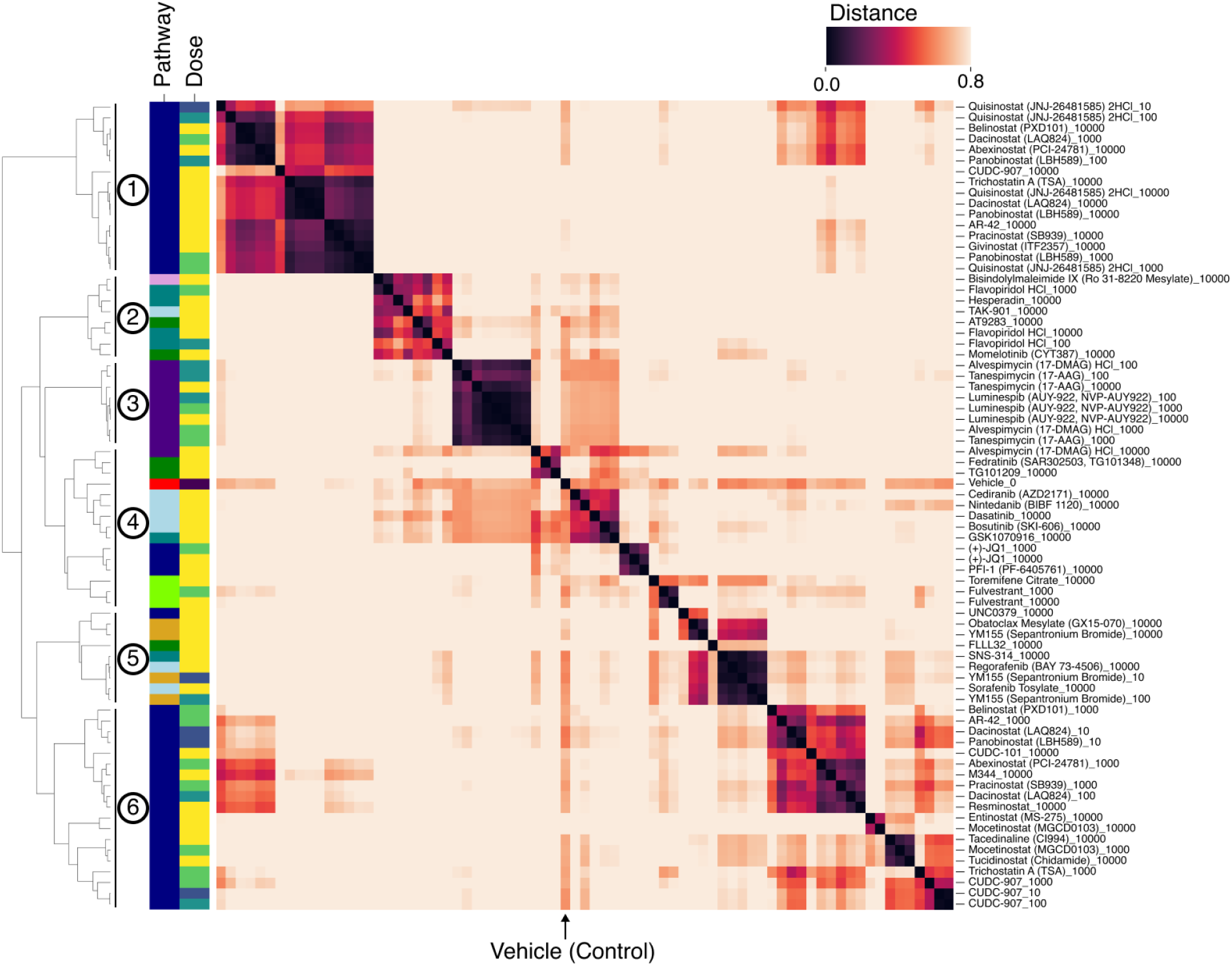
Copy of Figure S14e with row labels for each drug-dose combination.

**Figure S17:**
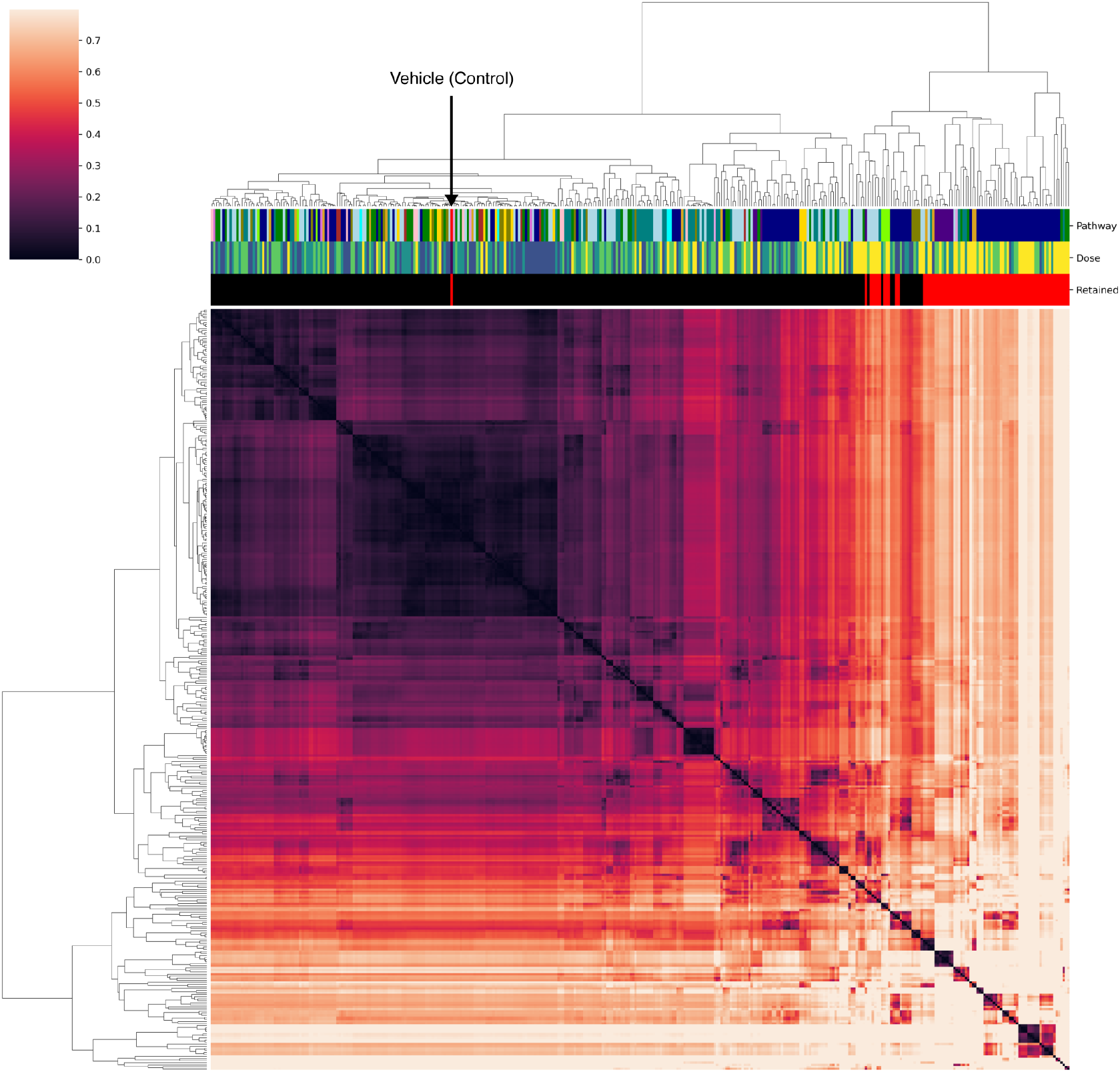
Sample distance matrix of all 369 samples (92 drugs at four doses and vehicle) used in the analysis for the MCF7 cell line. The columns are annotated by each drug’s pathway annotation from the original study, dosage level, and whether the sample was retained for the remaining analysis (top 20 percent of samples based on distance from vehicle). The hierarchical clustering was performed with the Ward variance minimization algorithm.

**Figure S18:**
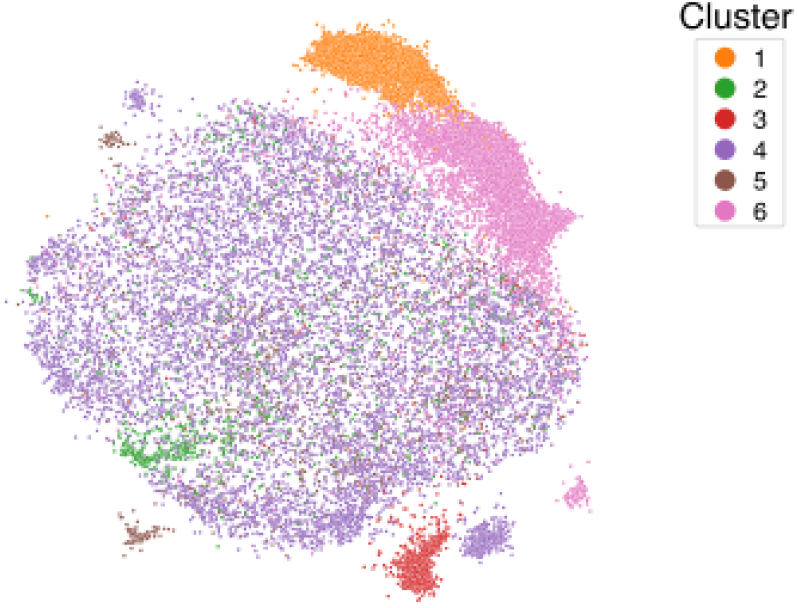
MDE of the *z* latent space from MrVI of data from the MCF7 cell line colored by Cluster as labeled in Figure S14e. Clusters ➅ and ➀ capture a large group of samples with similarly divergent cell states corresponding to HDAC inhibitors.

**Figure S19:**
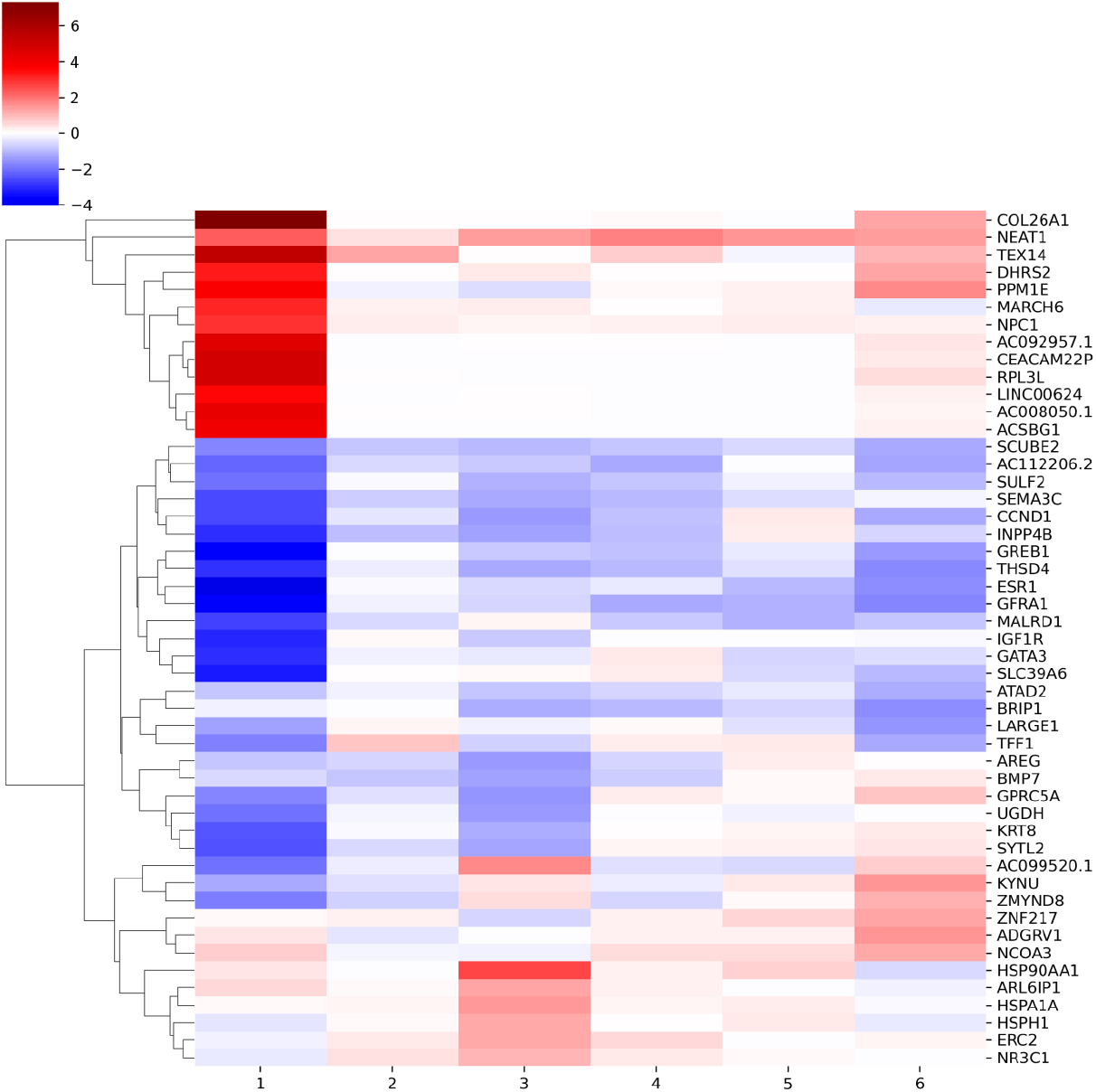
Heatmap of LFCs of the top DE genes identified by MrVI in the MCF7 cell line data. Each column corresponds to a cluster as labeled in Figure S14e. Displayed is the LFC estimated by MrVI averaged over all cells. Concordant with the distance matrix, we find the most significant and widespread gene-specific effects in cluster ➀.

**Figure S20:**
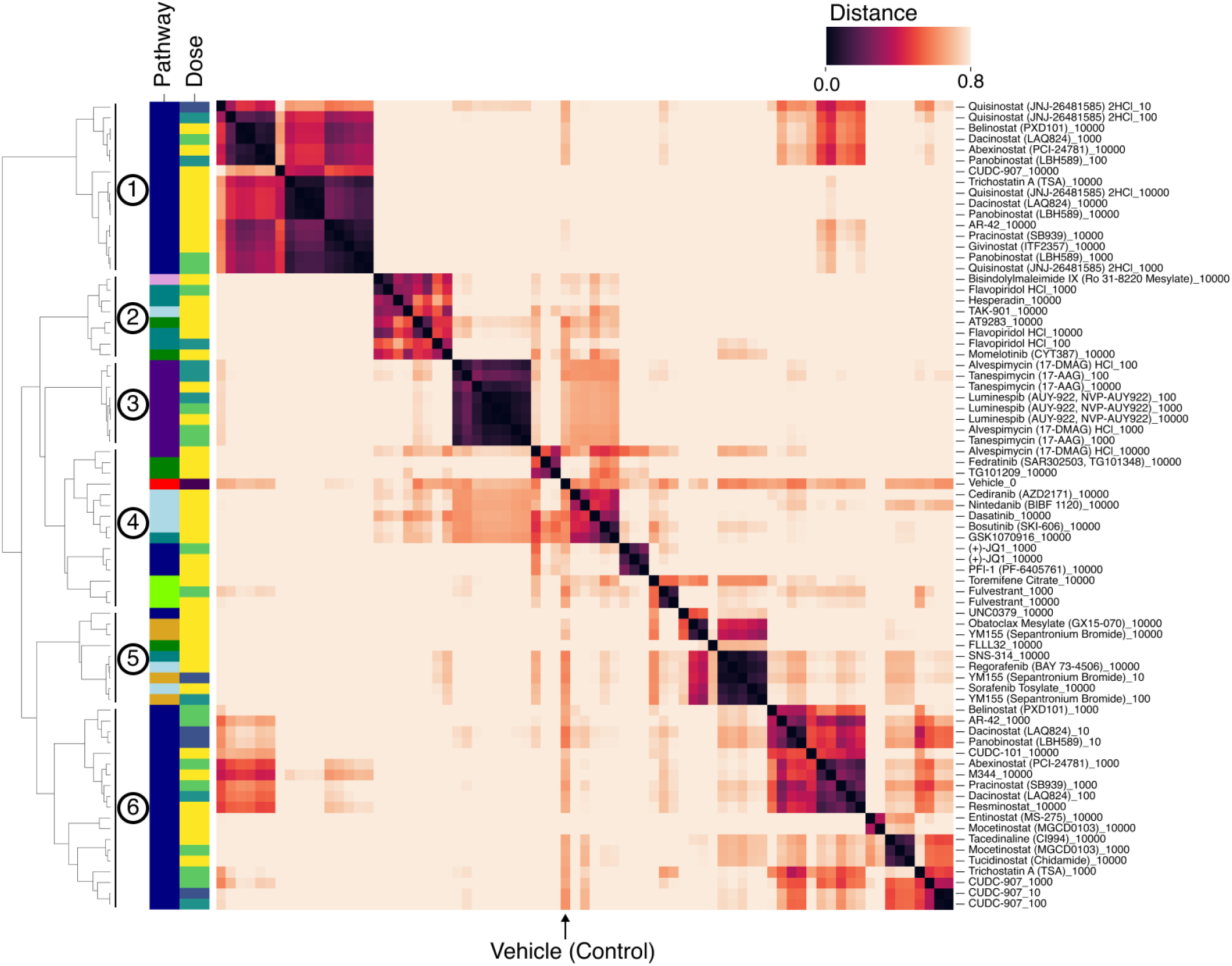
Copy of Figure S15e with row labels for each drug-dose combination.

**Figure S21:**
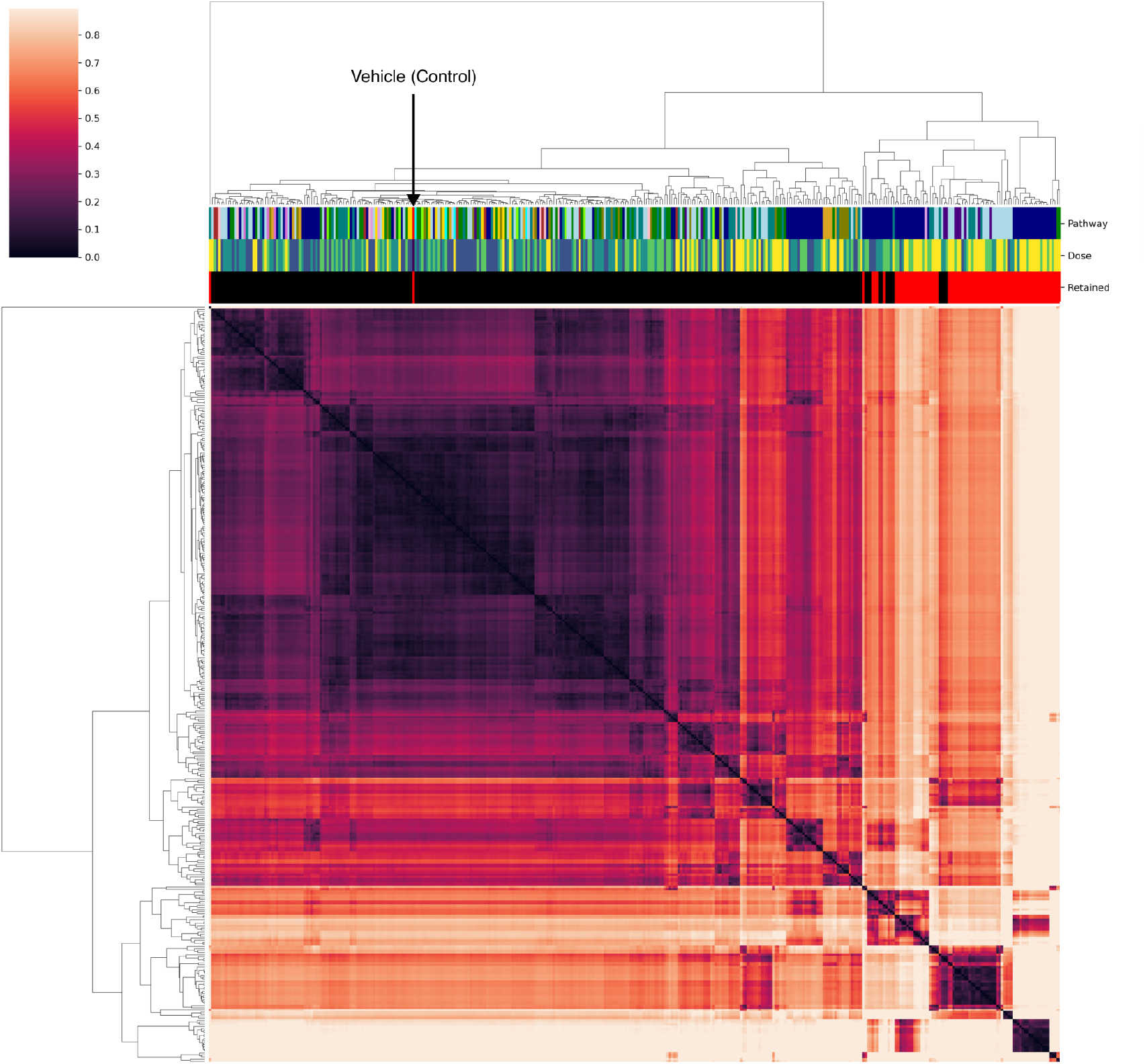
Sample distance matrix of all 369 samples (92 drugs at four doses and vehicle) used in the analysis for the K562 cell line. The columns are annotated by each drug’s pathway annotation from the original study, dosage level, and whether the sample was retained for the remaining analysis (top 20 percent of samples based on distance from vehicle). The hierarchical clustering was performed with the Ward variance minimization algorithm.

**Figure S22:**
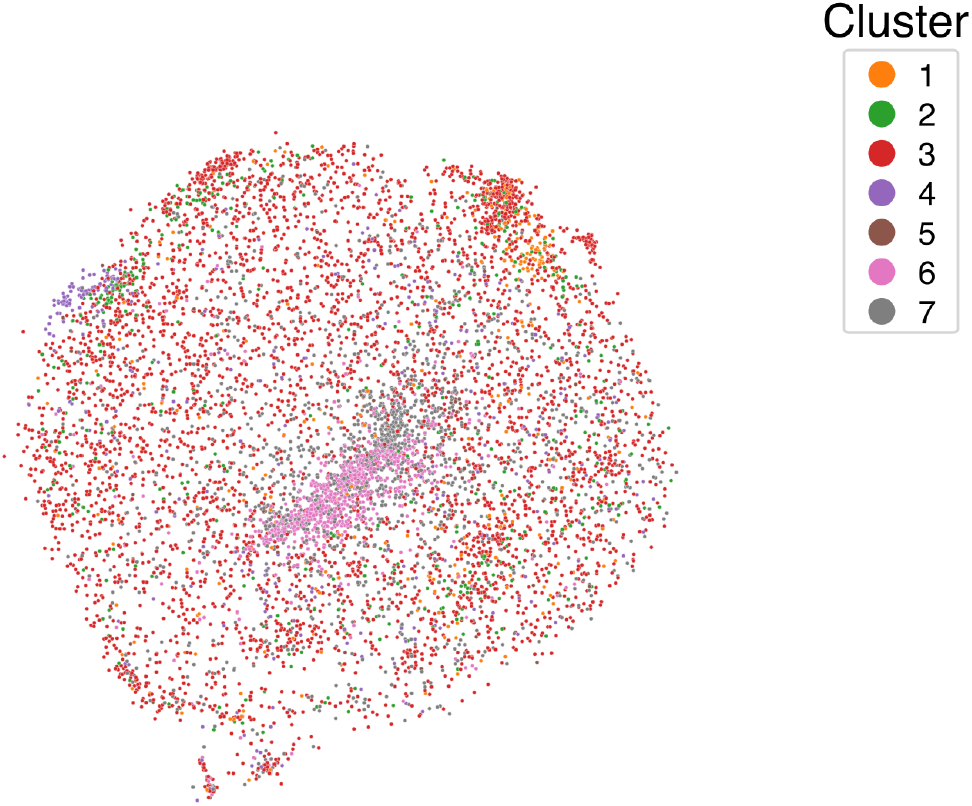
MDE of the *z* latent space from MrVI of data from the K562 cell line colored by Cluster as labeled in Figure S15e. We note that clusters ➅ and ➆ capture a large group of samples with similar transcriptomic responses corresponding to HDAC inhibitors.

**Figure S23:**
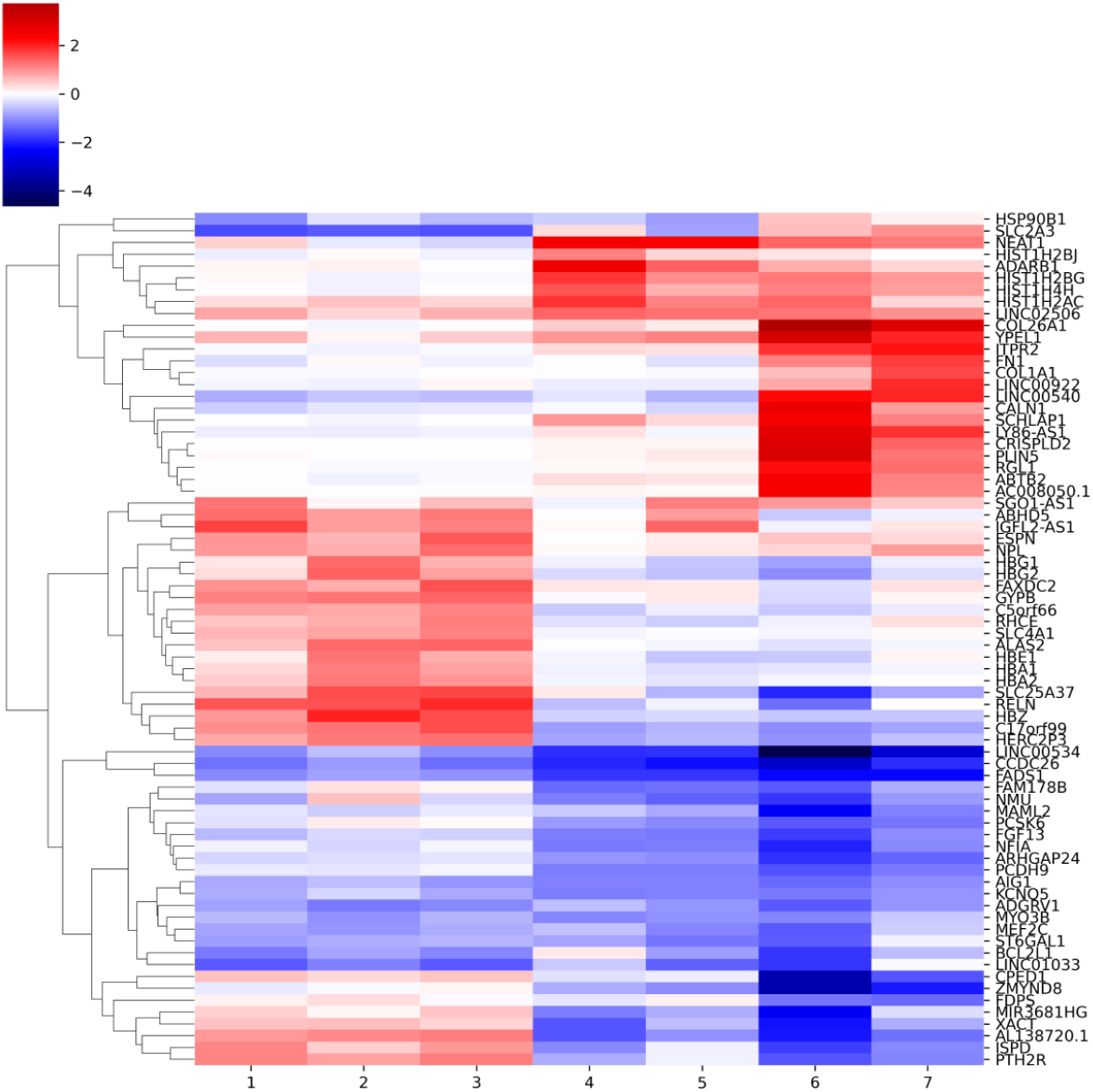
Heatmap of LFCs of the top DE genes according to MrVI over the K562 cell line data. Each column corresponds to a cluster as labeled in Figure S15e. Displayed is the LFC estimated by MrVI averaged over all cells. Concordant with the distance matrix, we find the most significant and widespread gene-specific effects in clusters ➅ and ➆.

**Figure S24:**
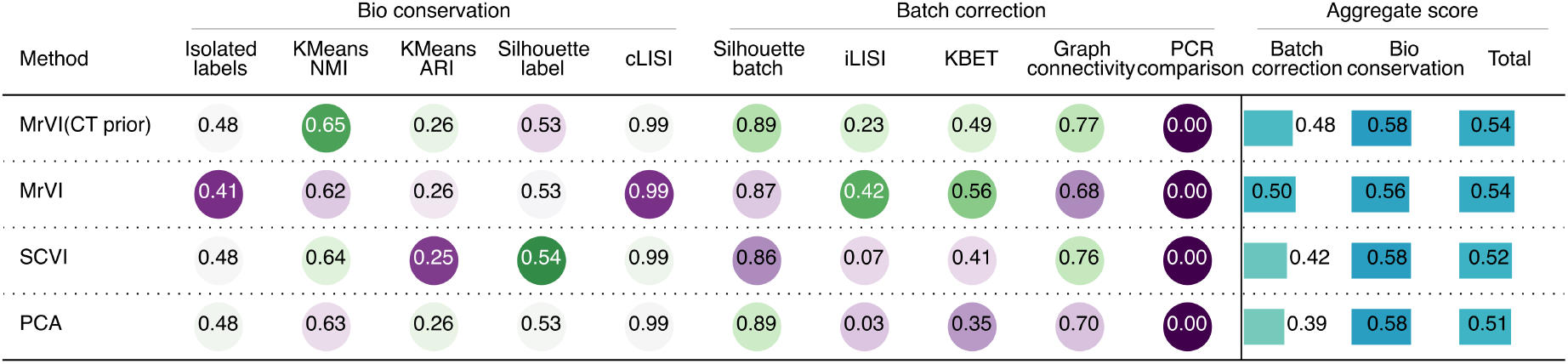
SCIB metrics computed on the IBD dataset. Tissue (colon or ileum) was used as batch key, and the original study’s annotations were used as cell-type labels as displayed in S27. MrVI (CT prior) uses the cell-type specific bias for the mixture weights using the existing annotations (**Methods**), while MrVI is the default version relying on mixture of Gaussians without cell-type biases.

**Figure S25:**
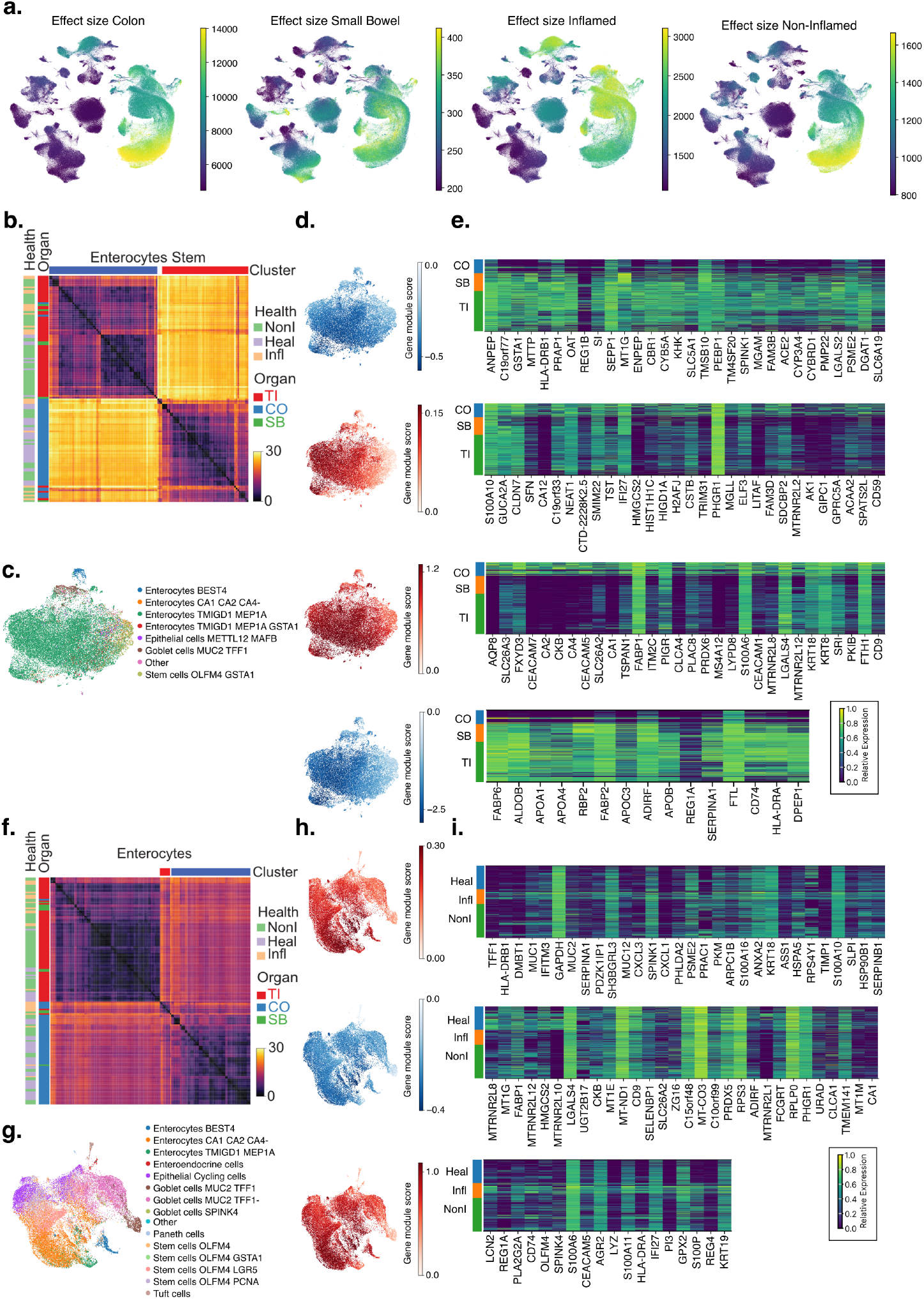
Unbiased analysis of Crohn’s disease dataset. **a**. We perform a guided analysis of differential gene expression (**Methods**). We compute the effect strength by computing the L2 norm of the covariate-specific vector in the z representation for each of the covariates (**Methods**). The effect size for the colon and small bowel are computed with ileum defined reference tissue, while the effect size for inflamed and non-inflamed biopsies is calculated with healthy biopsies as reference. **b**. Sample distance matrix of all cells annotated by us as *Enterocytes Stem* (immature epithelial cells). We find a separation into two distinct groups that correspond to the ileum (TI) / small bowel (SB) and the colon (CO) “Organ”. We find no strong subgrouping of samples with different disease status (Health in legend, NonI = non-inflamed, Heal = healthy, Infl = inflamed). **C** Original cell-type labels overlaid with a UMAP embedding based on MrVI embedding of all cells labeled as *Enterocyte Stem* cells. **d**. Scores of four gene modules identified between both highlighted clusters in the sample d matrix overlaid on UMAP plot. The score is the signed sum of all up- or downregulated LFCs within a module. **e**. The raw gene expression after library-size normalization and log1p-transformation is displayed in the heatmap and the heatmap is stratified by “Organ”. All genes within the respective module are displayed. The heatmap is grouped by the tissue. Expression values are displayed after standardizing gene expression values from 0 to 1 for each gene. **f-i** Same analysis as in **b-e** for *Enterocytes*. The samples show stronger disease stratification, revealing a third distinct cluster of inflamed samples. (h.) Gene modules show modules upregulated in inflamed biopsies in red, while the down-regulated modules are shown in blue.

**Figure S26:**
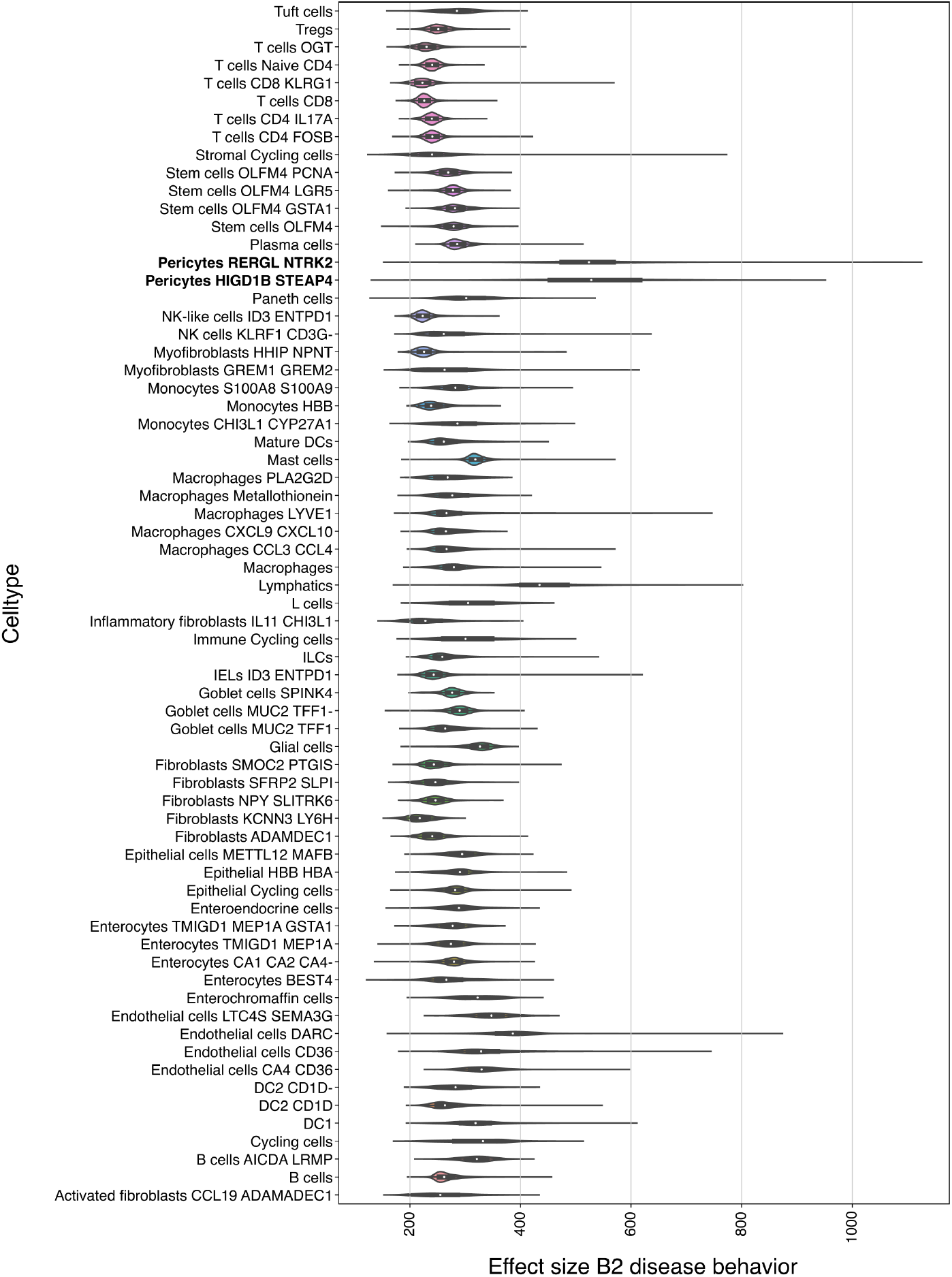
Violin plots of effect sizes of B2 disease behavior split by original cell-type labels. We quantify the estimated effect size displayed in **Figure 5a right**. Displayed is the violin plot for each cell type, and the plotted values are the estimated effect size for each cell. Highlighted in **bold** are both pericyte cell types that have the highest effect sizes.

**Figure S27:**
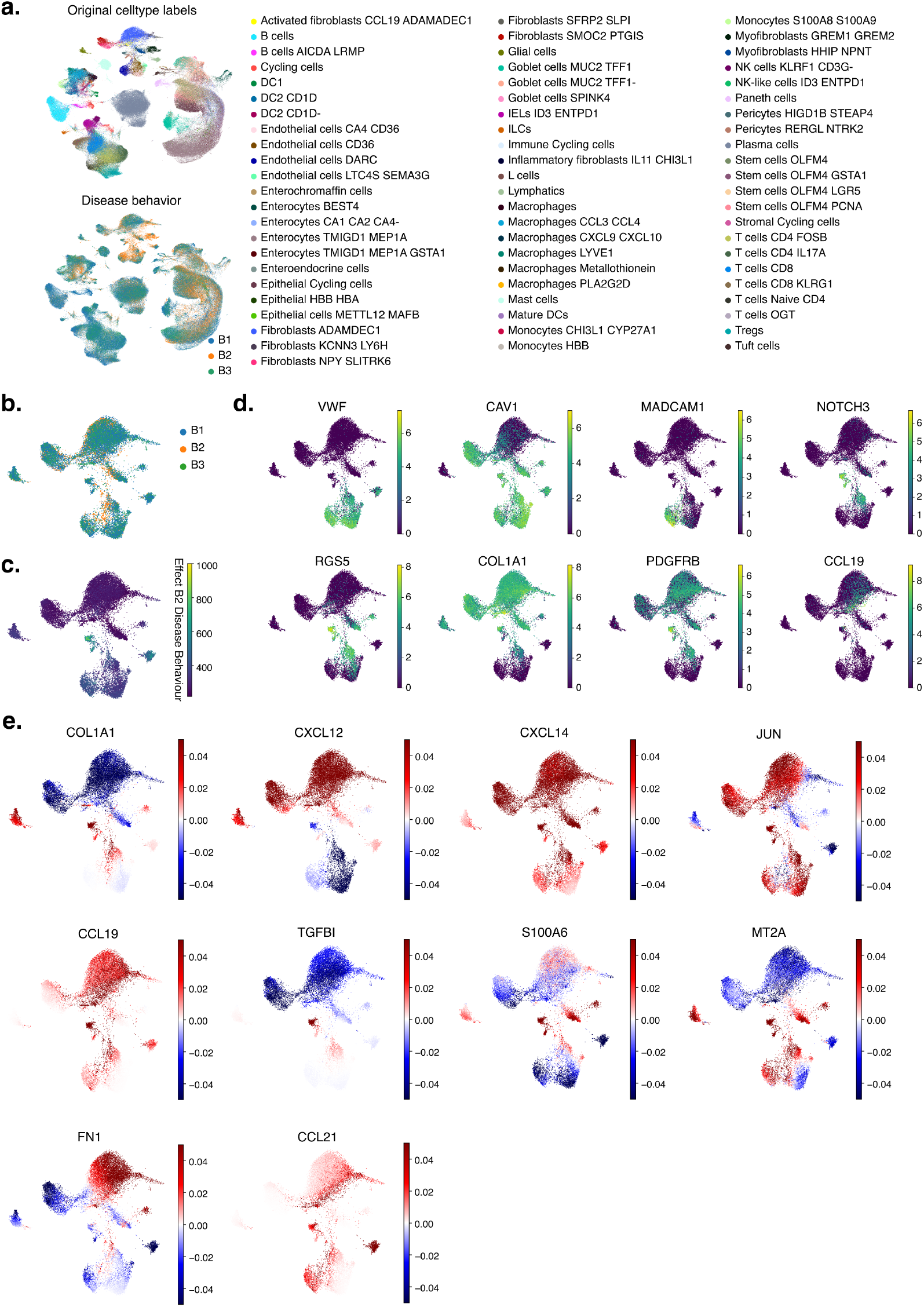
Additional analysis of stromal cells in stenosis. **a**. UMAP colored by all cell types in the original study as well as colored by disease behavior. **b-e** UMAP subset to stromal cells. **b**. colored by disease behavior. **c**. Same display as in **Figure 5a** of effect size of disease behavior B2 highlighting high score in a subset of pericytes. **d**. Raw gene expression after library-size normalization and log1p-transformation for marker genes of endothelial-to-mesenchymal transition. **e**. Additional predicted LFCs in patients with stenosing course of disease (B2) versus patients with B1 disease behavior patients estimated by MrVI.

